# Hybrid Neural Networks of the Olfactory Learning Center in the *Drosophila* Brain

**DOI:** 10.1101/2023.12.06.570338

**Authors:** Li-Shan Cheng, Ching-Che Charng, Ruei-Huang Chen, Kuan-Lin Feng, Ann-Shyn Chiang, Chung-Chuan Lo, Ting-Kuo Lee

**Author notes:** These authors contributed equally (Li-Shan Cheng, Ching-Che Charng). These co-corresponding authors contributed equally (Ann-Shyn Chiang, Chung-Chuan Lo, Ting-Kuo Lee).

## Abstract

Biological signal encoding is shaped by the underlying neural circuitry. In *Drosophila melanogaster*, the mushroom body (MB) houses thousands of Kenyon cells (KCs) that process olfactory signals from hundreds of projection neurons (PNs). Previous studies debated the connectivity between PNs and KCs (random vs. structured). Our multiscale analysis of electron microscopic data revealed a hybrid network with diverse synaptic connection preferences and input divergence across different KC classes. Using MB connectome data, our simulation model, validated via functional imaging, accurately predicted distinct chemical sensitivities in the major KC classes. The model suggests that the hybrid network excels in detecting food odors while maintaining precise odor discrimination in different KC classes. These findings underscore the computational advantages of this hybrid network.

## Introduction

Olfaction is crucial for animal survival and reproductive success, demanding efficient differentiation between food sources and irrelevant odors that translate into innate and learned behavioral responses. In the case of a fruit fly, *Drosophila melanogaster*, the MB governs olfactory associative learning (*1, 2*). This process commences with compressing olfactory signals from a multidimensional chemical space into a streamlined functional coding, orchestrated by approximately 52 glomeruli within the antennal lobe (AL) (*3-6*). These signals then diverge into projections, transmitting olfactory information from around 100 monoglomerular PNs across the 52 glomeruli to an assembly of roughly 2000 KCs (*7-9*). These olfactory cues subsequently acquire functional significance through altering connection weights between sparsely active KCs and output neurons (MBONs) and thereby triggering actions through MBONs (*10-12*). Through processing in the triadic PN-KC-MBON hierarchical network, fruit flies distinguish distinct chemical compounds, generalize experiences to akin odors, and adeptly perceive their environment (*13, 14*). However, the intricate neural circuitry translating odor representation from the AL glomeruli to KC coding underpinning the impressive functionality of the fruit fly olfactory system remains an enigma (*15, 16*).

Earlier investigations introduced two contrasting hypotheses concerning PN-to-KC connectivity: random versus structured (*17-28*). The random hypothesis posits random PN-to-KC connections, backed by functional response variations and a lack of precise cellular-level circuit specification (*20, 24*). A specific type of KC exhibited varied reaction profiles to the same odor across different brains(*24*). For instance, Caron et al. (2013) (*20*) labeled inputs of 200 KCs through dye injection, discovering that no two KCs shared the same glomerular input combination. The spatial patterns of activated KCs displayed inconsistencies among animals, implying a random architecture (*24*). However, this seemingly random connectivity could arise from individual network rewiring (*29*). Furthermore, MBON activity associated with valence encoding exhibited higher inter-individual correlation than expected by chance (*27*). The manner by which the calyx’s seemingly random connectivity produces consistent MBON responses without the learning-related protein Rutabaga remains obscure (*11*).

In contrast, the structured hypothesis posits a degree of preferential connectivity between PNs and KCs at a population level, an arrangement maintained across individuals (*17, 18*). This hypothesis finds support in various observations: (i) Functional imaging demonstrated consistent spatial calcium response patterns in KCs across diverse individuals for the same odor (*25*). (ii) Anatomical studies revealed significant overlap between specific glomerular PN axon arbors and dendritic arbors of particular KC classes (*17, 28*). (iii) Neuronal spatial innervation of PNs could be clustered based on a metric assessing neuronal distance (*30*). (iv) Mutant flies without normal olfactory co-receptors displayed comparable glomerular input ratios to KC classes across diverse individuals (*21*), implying genetically determined PN-to-KC connectivity preferences.

Early studies were hindered by low image resolution and incomplete sampling, yielding partial connectome reconstructions (*20*). However, the recently released FlyEM database offers nearly comprehensive synaptic-level connectome data, comprising 106 monoglomerular PNs and 1745 KCs (*31*). This heightened connectivity insight permits revisiting whether the calyx harbors concealed wiring preferences and how this structure influences olfactory information processing. Connectomics analysis unveiled three distinct morphological types of PN boutons containing varying presynaptic vesicles (*32*), while food-responsive PNs exhibited a community structure connecting to KCs beyond a random bouton model (*22*). The heterogeneity of PN boutons prompted deeper investigation into the inherently structured nature of the calyx’s connectome and its contribution to the MB’s olfactory signal encoding for heightened odor perception.

Here, we capitalize on accessible connectome data to quantify synaptic connectivity preferences between glomerulus-specific PNs and three major KC classes: γ, α’/β’, and α/β (*33*). Employing multi-scale analyses, we reveal connection preferences at various levels, ranging from synapses to populations. Specifically, PN cluster-to-KC class connections demonstrate a hybrid connectivity pattern marked by differing preference degrees with distinct convergent or divergent patterns. We validate this hybrid connection scheme through computer simulations of KC response profiles to diverse odors, comparing results with *in vivo* calcium imaging. Our computer model further illustrates the potential advantages of this network in olfactory coding. Ultimately, our study illuminates intricate connectivity patterns and functional interrelationships within the PN-to-KC hybrid network, shedding light on mechanisms guiding odor processing and response generation within the insect brain.

## Results

### Diverse Preferences in PN-to-KC Connections

To unravel potential connection patterns of the PN-to-KC network within the calyx (fig. S1A and B), we commenced by dissecting connectivity using the comprehensive FlyEM dataset (*31*) (Fig. 1A), an electron microscopy-based synaptic connectome derived from a female fly. Our examination of connection preferences (G) encompassing approximately 100 monoglomerular PNs related to olfaction and three distinct KC classes (γ, α’/β’, and α/β, as illustrated in Fig. 1B) entailed a comparison of the real network with randomly shuffled networks (for methodological details, refer to Methods). Since each glomerulus PN and its boutons demonstrate disparate connection capacity, our shuffling algorithm preserved the connection number between individual PNs and KCs, maintaining the heterogeneous counts of KCs connected to a given PN’s bouton across different PN types (fig. S1, C to F). The results of our analysis brought to light that each KC class exhibits preferred connections stemming from a specific subset of PNs (Fig. 1, B and C, and fig. S2). Specifically, the γ class features preferred (G_γ_ > 2) or disfavored (G_γ_ < -2) connections with 33 glomeruli (∼ 63%), the α’/β’ class features preferred (G_α’/β’_ > 2) or disfavored (G_α’/β’_ < -2) connections with 19 glomeruli (∼36%), and the α/β class features preferred (G_α/β_ > 2) or disfavored (G_α/β_ < -2) connections with 39 glomeruli (∼75%) (Fig. 1B). Moreover, we observed a strong negative correlation between the preference score distributions of the γ and α/β classes (Pearson’s correlation coefficient = -0.96, p < 0.001). A similar trend emerged when considering connection weightings (synapse numbers) (fig. S3). In contrast, after shuffling, the heterogeneous connection weights among PN clusters and KC classes become homogeneous (fig. S2).

**Fig. 1.**
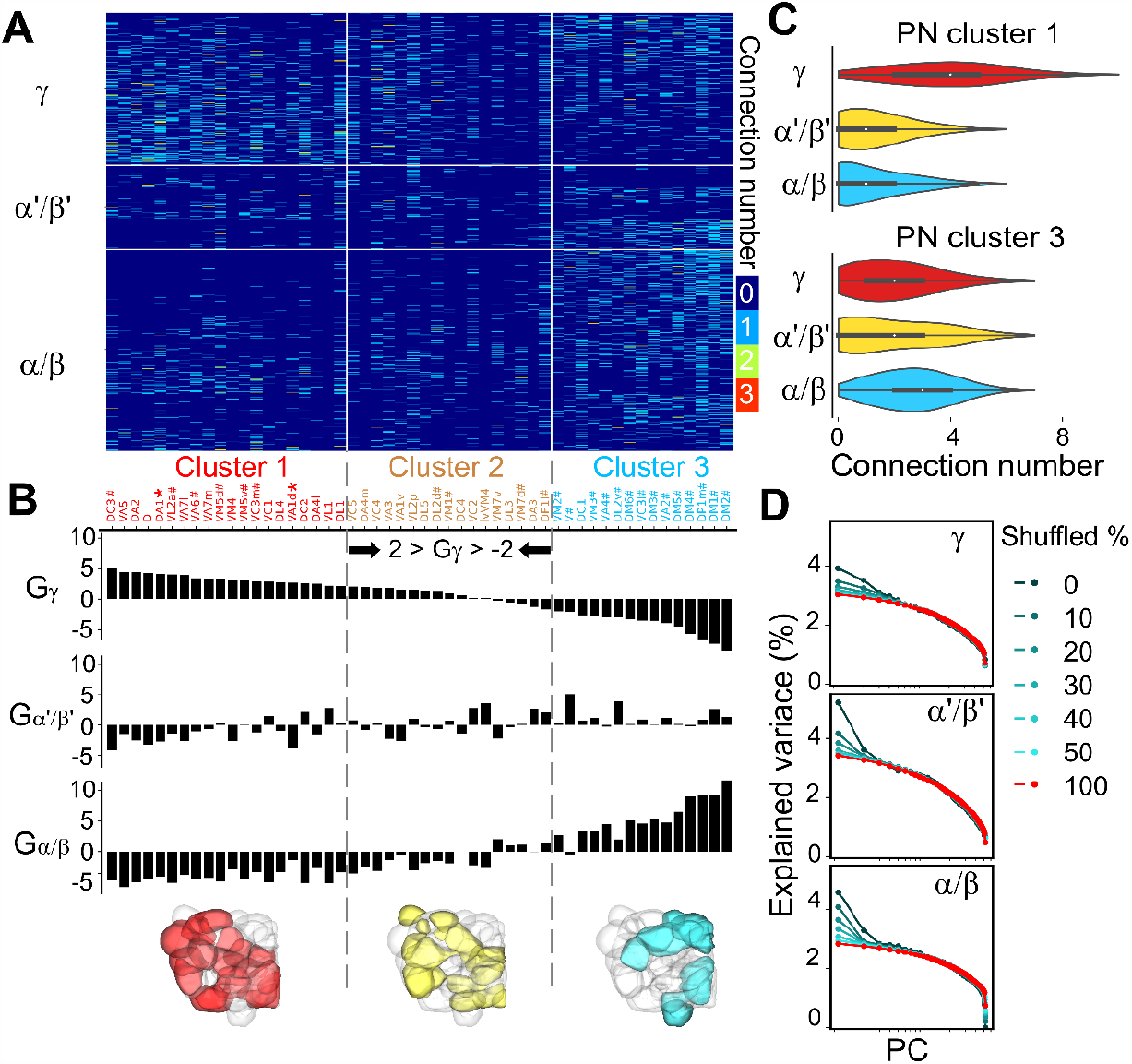
Preferential connectivity between olfactory PNs and three classes of KCs. (**A**) The connectivity matrix of PN-to-KC synapses derived from the FlyEM dataset (*31*). PNs originating from individual glomeruli were categorized into three clusters based on their connection preferences (see **B**). The color coding indicates the number of monoglomerular PNs connected to a particular KC. (**B**) The connection preferences indicated by the G_i_ score between a KC class i (i = γ, α’/β’, or α/β) and PNs from a particular glomerulus (see Methods). The glomeruli are ordered by the G_γ_ score. Dashed lines indicate the range of G_γ_ scores of the shuffled results between -2 and 2. # indicates food-responsive glomeruli (*22*). * indicates pheromone-attractive glomeruli for a female fly (*66*). (**C**) The frequency distribution of PN numbers received by each individual neuron in three KC classes (see more details in fig. S2). (**D**) Analysis of global input preferences revealed by PCA on the PN-to-KC connection matrix after global shuffling of upstream PNs for individual KCs within the same class (refer to fig. S5 for detailed information). The color indicates the shuffled ratio of each KC class.

Next, we classified the 52 glomeruli into three distinct clusters based on the preference score G_γ_ (cluster 1: G_γ_ ≥ 2, cluster 2: 2 > G_γ_ > -2, cluster 3: G_γ_ ≤ -2, as depicted in Fig. 1B). Importantly, we found that glomeruli within the same cluster shared analogous functionality, with an astounding 93% of cluster 3 glomeruli (labeled by “#”) displaying responsiveness to food-related odors (*22*) (Fig. 1B).

Furthermore, we probed whether KC subclasses also displayed connection preferences originating from different glomeruli. Employing the same analytical approach, we uncovered that such connection preferences were indeed manifest in several subclasses (fig. S4). Specifically, the α’/β’-ap, α/β-p, α/β-s and γ-d subclasses emerged as recipients of inputs from certain cluster 2 glomeruli, signifying their distinctive odorant representation, while the α’/β’-m subclass predominantly integrated food-related information similar to the α/β class.

### Global Input Preferences of different KC Classes

We proceeded to establish input preferences by contrasting Principal Component Analysis (PCA) results between the observed network and networks with shuffled connections. The quantification of global input preference involved calculating the level of randomness after global shuffling of connections within the calyx (see Methods), regardless of their original spatial locations.

We executed separate shuffling procedures for each of the three KC classes, varying proportions from 0% to 100% (Fig. 1D and fig. S5). When a larger contingent of KCs receives comparable glomerular input combinations compared to the expectation from random chance, the first principal component (PC1) of the PN-to-KC network displays heightened variance compared to the entirely shuffled network. Our findings disclosed diverse degrees of preference across KC classes. For instance, a shuffling with ratios of 45% for γ, 45% for α’/β’, and 65% for α/β demonstrated comparable variance with fully shuffled groups (Fig. 1D and fig. S5). These observations intimate a “hybrid” network structure within the calyx, encompassing the γ network, which shows relatively modest input preference but still gathers greater input from PN cluster 1 (Fig. 1c and fig. S2). In contrast, the α/β network displays more pronounced input preference and mainly receives odor information from PN cluster 3 (fig. S2). This hybrid network configuration potentially empowers a fly’s brain to encode diverse odor types (see Discussion).

Discrepancies between our findings and prior research (*20*) may derive from two factors: (1) the scale of sampling, encompassing both neuron and connection numbers, and (2) a plausible sampling bias toward specific KC classes. To ascertain the minimum number of neurons essential for detecting input preference, we conducted random subsets sampling spanning from 5% to 100% of KCs, calculating the mean variance of the first principal component. Our results underscore that to unveil the distinction between a genuine network and shuffled sub-networks, at least 100 KCs are needed for α’/β’ and γ, while 80 KCs suffice for α/β (fig. S6). Furthermore, detecting input preference becomes more intricate when connection data for a particular KC class remains incomplete. Even with full γ KC collection and only 50% of connections, input preference wasn’t discerned for γ. When scrutinized with the same sampling count utilized by Caron et al., (*20*) the subsampled data consistently displayed random-like traits (fig. S7). Hence, the inference of a randomly structured calyx network may find an explanation in inadequate sampling sizes.

### Spatial Segregation of PN Axons and KC Dendrites

Next, we examined whether the observed connection preferences could be attributed to the spatial positioning of PNs (Projection Neurons) and KCs (Kenyon Cells) within the calyx. This investigation was conducted across three levels: neurite, synapse, and bouton/claw. An examination of the traced skeletons of all calycal neurites stemming from PNs revealed that a majority of cluster 3 PNs tended to congregate in the ventral-posterior region. In contrast, cluster 1 PNs exhibited a tendency to cluster more prominently in the dorsal-anterior domain (Fig. 2A and fig. S8). Regarding the distribution of dendritic arbors of the three KC classes within the calyx, the α/β and α’/β’ classes were densely concentrated in the ventral-posterior region, while the γ class displayed a more uniform distribution (Fig. 2B and fig. S8).

**Fig. 2.**
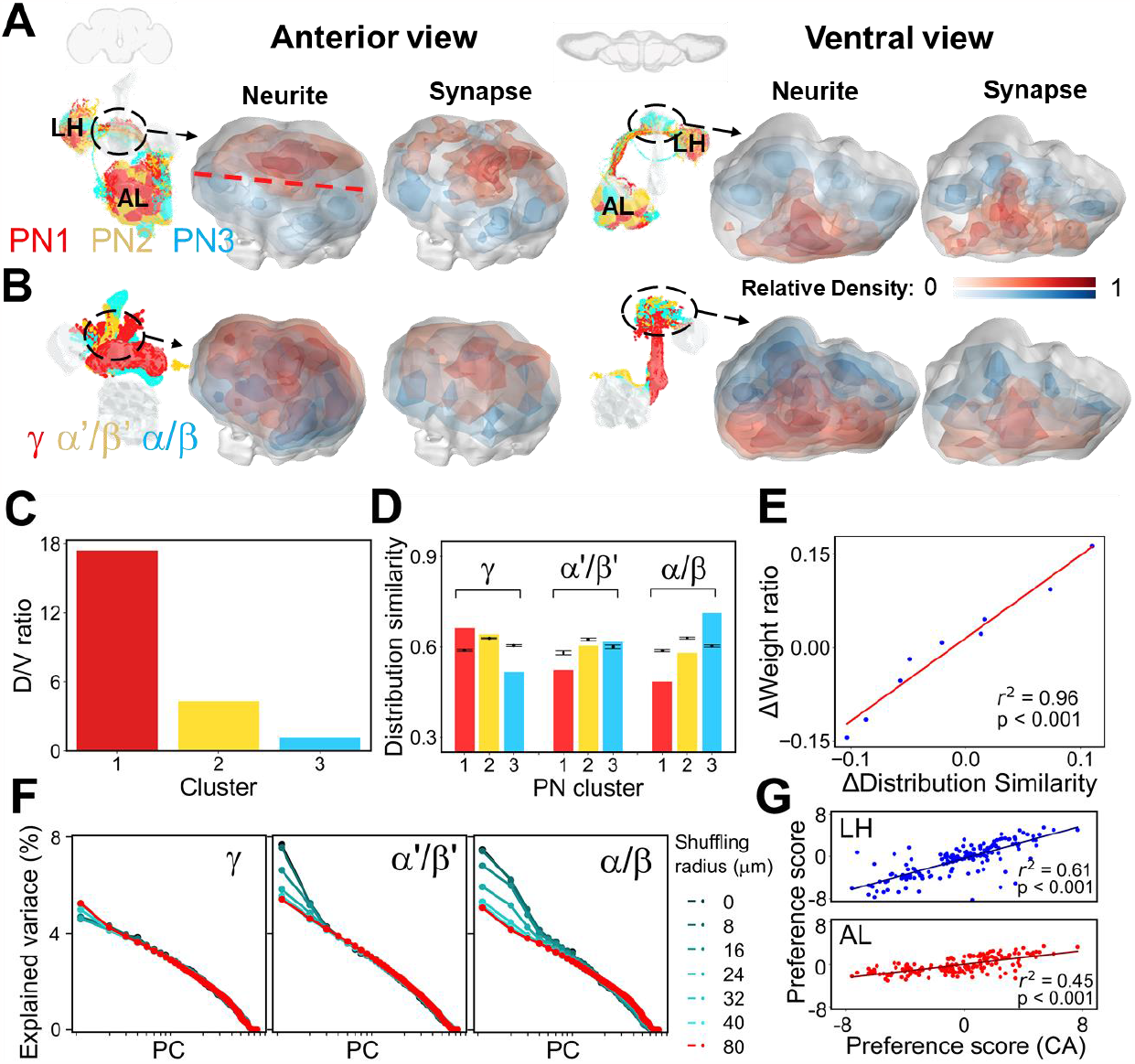
Preferential spatial connectivity between PN clusters and KC classes. (**A**-**B**) Spatial distribution of neurites and PN-to-KC synapses from each PN cluster (**A**) and KC class (**B**) shown by color code. AL: antennal lobe, LH: lateral horn, CA: calyx. (**C**) Quantitative spatial distribution of PN neurites. The dorsal/ventral (D/V) compartments are separated by the red dashed line in (**A**). (**D**) Distribution similarity between presynapse and postsynapse in PN-to-KC connections compared with shuffled data (error bars: ± s.d.). (**E**) Correlation between the distribution similarity of PN-to-KC synapses and their input connection weight ratio (see Methods, fig. S2E) quantified by the difference between the real network and shuffled networks, as shown by the linear regression plot (r^2^ = 0.96, p < 0.001). (**F**) Analysis of local input preferences quantified by PCA after local random shuffling of KC claw inputs. Colors indicate different shuffling radii (refer to fig. S15 for detailed information). (**G**) Correlation of spatial innervation preference between each glomerulus and each PN cluster by the comparison between the real network and shuffled networks (see Methods), as shown by the linear regression plot (for the comparison between CA and LH, r^2^ = 0.61, p < 0.001; for the comparison between CA and AL, r^2^ = 0.45, p < 0.001).

To assess the statistical significance of these distributions, we conducted a categorical shuffling test. This test involved permuting the categorical classification of both PNs and KCs to determine their deviation from random placement in the calyx (fig. S9 and fig. S10, see Methods for more details). Our results indicated that cluster 1 PNs and cluster 3 PNs occupied significantly different areas compared to the shuffled results (fig. S9A). Similarly, the distributions of α/β and γ classes also showed notable deviations from randomness (fig. S9C).

To investigate whether PN clusters exhibited a preference for specific regions in their neurite projections, we conducted a shuffling of KC identities. Our findings suggested that PN cluster 1 tended to project toward areas populated by more γ KCs, while PN cluster 3 projected toward the dendritic arbors of α’/β’ and α/β KCs (fig. S9E and fig. S10A). A more detailed examination of glomerular projection patterns revealed the segregation of cluster 1 PNs and cluster 3 PNs, not only within the calyx but also in the lateral horn (LH) (fig. S11).

### Distinct Spatial Organization of PN-to-KC Synapses

We then proceeded to scrutinize the population-level spatial distribution preferences of synapses for PN clusters and KC classes (Fig. 2, A and B, fig. S8, and fig. S9). Similarly, we performed a categorical shuffling of KC synapses to juxtapose the observed distribution with a random counterpart, while keeping the connection number for each neuron constant (Fig. 1). Our results revealed that cluster 1 PNs exhibit a greater synaptic overlap with γ, while α/β and α’/β’ manifest higher congruence with cluster 3 PNs (Fig. 2D, and fig. S9F).

Next, we calculated the correlation between spatial distribution similarity and the connection weight ratio (see Methods). To investigate this relationship, we employed a method involving shuffling KC synapses and comparing the observed network’s quantities from FlyEM^31^ with those obtained from the shuffled versions. Notably, we found a strong correlation between these two quantities (Fig. 2E, with r^2^ = 0.96 and p < 0.001). This alignment corresponds with the predictions of Peter’s rule (*34, 35*), indicating that if axonal and dendritic arbors exhibit greater overlap, neurons could forge more synaptic connections. Moreover, when we arranged glomeruli based on spatial innervation preference for γ KCs, we observed patterns akin to those achieved through connection preference-based sorting (Fig. 1B) across all three tiers (fig. S10). Furthermore, the synaptic spatial innervation preferences between glomerular PNs and subclasses of KCs (fig. S12) also show a comparable trend to the connection preference (fig. S4). The spatial innervation preferences held a robust correlation with the connection preferences (fig. S13, with r^2^ = 0.857 and p < 0.001), highlighting that connection preferences can be attributed to spatial innervation preferences.

Given that the FlyEM dataset (*31*) originated from an individual fly, we extended our examination of PN cluster spatial distribution to another EM dataset, the FAFB dataset (*36*). We observed consistent patterns (fig. S14), indicating the presence of spatial innervation preferences across diverse individuals.

### Local Input Preferences of KC Network

The non-random spatial distribution of PN-to-KC connections led us to undertake a more detailed investigation of the local organization. Our aim was to determine whether there is a specific preference for connectivity between the axonal boutons of PNs and the dendritic claws of KCs within the calycal microglomeruli. Considering the significant energy requirements associated with neurite branching (*37*), it is plausible that KC claws exhibit a tendency to form connections primarily with adjacent PN boutons. To facilitate this exploration, we devised a local shuffling algorithm (see Methods), allowing for localized randomization of connections within the calyx. This algorithm permits claws to interchange their upstream boutons, solely if the relative distance is shorter than a certain threshold *R* (fig. S15). Remarkably, within a 40 μm radius, the first principal component (reflecting connectivity variance) witnessed a 30% decrease for α’/β’ and a 50% decrease for α/β, respectively (Fig. 2F and fig. S15). In contrast, the principal component strengths of γ presented a more uniform distribution (Fig. 2F). These outcomes underscore that the local distribution of claws is intimately linked to the stereotypical formation of α’/β’ and α/β, but this isn’t as prominent for γ. Consequently, this convergent structure appears to stabilize odor representation, particularly for food-related odors. Our extensive investigation has unveiled a panorama of spatial connectivity preferences, spanning from the neurite to the synapse level.

### Segregation of PN Terminals across AL, calyx and LH

To further explore whether the spatial connectivity preferences we observed within the calyx extended to two other related neuropils, we generated density maps of the three PN clusters in the AL and the LH based on their total postsynaptic and presynaptic coordinates, respectively. Notably, our analysis of the postsynaptic density map revealed that cluster 3 PNs predominantly occupy the middle glomeruli, while cluster 1 PNs exhibit a distribution across the lateral glomeruli (Fig. 2A and fig. S16, A and B). Within the LH, the presynaptic density maps for cluster 3 PNs coalesce at the center, whereas cluster 1 PNs display a bilateral distribution pattern (Fig. 2A and fig. S16, A and B).

When comparing the spatial innervation preferences at the synaptic level, we observed that cluster 1 and cluster 3 PNs exhibited distinct spatial distributions from each other in both the LH and AL (fig. S11 and fig. S16, C to F). An interesting observation emerged when we compared the preference scores for each glomerulus of PNs between cluster 1 and cluster 3 – an anti-correlation was also evident in both the AL and LH, consistent with what we observed in the calyx (fig. S16, E to G). This pattern of spatial connectivity preferences appears to be a consistent and intriguing feature of neural connectivity across these related neuropils.

Remarkably, a closer examination of the relationship between a glomerulus’ preferences for a specific PN cluster in both the calyx and LH using linear regression analysis revealed strong and consistent correlations (Fig. 2G and fig. S16H). These patterns were mirrored when comparing the calyx to the AL, suggesting a general phenomenon (Fig. 2G and fig. S16H).

Furthermore, it is worth noting that cluster 3 PNs not only displayed this connectivity pattern but were also found to receive inputs from the same type of sensilla (*38*) (fig. S17). This remarkable uniformity across various olfactory centers underscores the presence of an inherited wiring pattern within the calyx connectome. These findings hold the potential to provide valuable insights into the fundamental mechanisms governing olfactory encoding.

### Functional Correlates of Calycal Connection Preferences

To assess the functional implication of PN connection preferences within diverse olfactory information centers, we embarked on an inquiry to ascertain whether AL glomeruli within the same PN clusters displayed similar responses to odors bearing comparable chemical attributes. Conventionally, glomeruli adhere to the labeled-line architecture, wherein they receive input from distinct odorant receptor neuron (ORN) types, thus inheriting response profiles (*39*). Previous studies have revealed the clustered responsiveness of ORNs to aliphatic or aromatic (arom) odorants (*3, 40, 41*). In our investigation, we scrutinized ORN responses to pure odor stimuli sourced from the DoOR database (*42*). This enabled us to pinpoint the most favored ligand candidate for each ORN, indicative of the odor eliciting the most robust response. Surprisingly, our observations unveiled an augmented sensitivity of cluster 3 glomeruli to short-chain esters featuring one to two-carbon chains, hinting at an unexpected interplay between functionality and convergent glomerular architecture (Fig. 3A). In contrast, the most favored ligand candidates for glomeruli in clusters 1 and 2 didn’t exhibit clear chemical resemblances. Instead, clusters 1 and 2 manifested broad responsiveness to alcohols, aromatics, esters, and aliphatics. These intriguing findings suggest that the PN-to-KC network’s connection preferences exhibit a pronounced association with the processing of olfactory information tied to specific functional groups, rather than encompassing a universal sensitivity to all foraging scenarios.

**Fig. 3.**
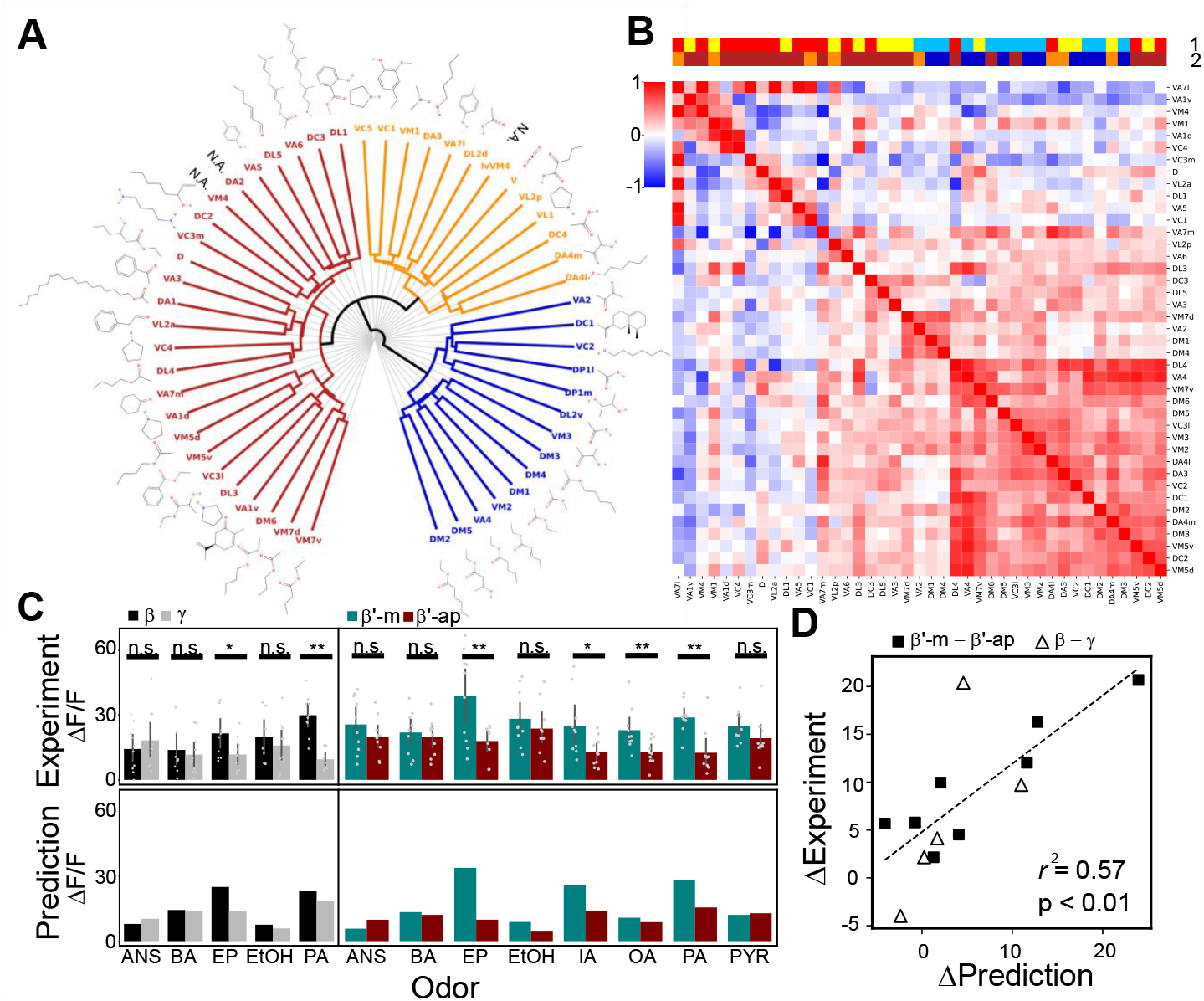
Preferential functional connectivity between PNs and KCs. (**A**) The most sensitive odorant for each glomerulus, based on the DoOR database (*42*). The glomeruli are classified into three clusters (indicated by different colors) based on their output correlations (see fig. S17). (**B**) The correlation matrix of ORN odorant responsive profiles. The color codes on the top two rows indicate (**1**) three preference clusters, as shown in Fig. 1C, and (**2**) glomerular output clusters, as shown in (**A**). (**C**) Experimental validation (upper panel) of predicted (lower panel) functional calcium responses (*Δ*F/F) to eight odorants (see Methods) at β, γ, β’-m and β’-ap MB lobes. *: p < 0.05, **: p < 0.01, and ***: p < 0.001 (Error bar = mean ± S.D., n =10 for each measurement, Wilcoxon Signed Rank Test). The predicted response is employed by numerical simulation using the DoOR database (*42*) and the FlyEM dataset (*31*) (see Methods). (**D**) Correlation between simulated and experimental calcium responses in the corresponding MB lobes. We quantify the response differences between β and γ lobes (triangles), and between β’-m and β’-ap (squares) for data from simulations and experiments in (**C**), as shown by the regression plot (r^2^ = 0.57, p < 0.01).

To further investigate the correlation in responsiveness between glomeruli, we delved into the composition of the top ten most responsive molecules of ORNs for each glomerulus. The outcomes indicated a heightened prevalence of esters within cluster 3 (fig. S18). Furthermore, we undertook a correlation analysis to quantify the likeness in ORN tuning for approximately 700 monomolecular odors. Remarkably, we identified a cluster of glomeruli demonstrating significantly high responsiveness similarity (Fig. 3B). Interestingly, these predominantly belonged to cluster 3, which converges toward overlapping downstream KCs (Fig. 3B and fig. S17). Conversely, the glomerular responsive profiles within both cluster 1 and cluster 2 exhibited relatively low intra-cluster correlation.

### Validating the Prediction of Odor Responses from Connectivity

Next, we combined functional data from the DoOR database (*42*) with connectivity insights from the FlyEM dataset (*31*) to validate our predictions about how different types of KCs respond to specific odors. Our simulations revealed distinct odor preferences among various KC classes: short-chain esters exhibited a preference for α/β and α’/β’-m classes (fig. S19). To confirm the accuracy of our predictive model, we conducted calcium imaging experiments using eight different odors, targeting specific regions within the MB lobes corresponding to different KC classes (fig. S20). The calcium responses to all tested odors closely matched our predictions, indicating consistent functional differences among MB lobes (Fig. 3C). Importantly, when we randomly shuffled the data in our simulation model, it failed to predict these functional disparities (fig. S21). Furthermore, we observed that these response differences between β and γ classes remained consistent across varying odor concentrations, ranging from 10^−6^ to 10^−2^ dilutions (fig. S22). This aligns with dose-dependent functional responses observed in ORNs (*40*) and PNs (*43*), as well as the spatial KC responses in the calyx (*25*) and memory-guided behaviors (*13*). The strong correlation between our experimental and simulation data was evident (r^2^ = 0.57, as shown in Fig. 3D). These findings collectively underscore the critical role of preferential connectivity between PNs and KCs in shaping sensory representation and conferring chemical sensitivity at the population level.

### Hybrid Network Architecture Diversifies Olfactory Coding Strategies

The validation of predictive odor responses strongly supports the idea of a hybrid network configuration, characterized by unique synaptic preferences and input divergence among different KC classes. To gain a deeper understanding of the functional significance inherent in this hybrid network, we performed neural network simulations using artificial odors to assess olfactory acuity and coding capacity (see Methods). In our evaluation of olfactory acuity, we carefully designed artificial odors by activating specific sets of glomeruli within clusters, guided by preference scores. We incorporated connection weights into our simulations, normalizing the weights based on the observed weight of a given KC relative to the total input PN connection weights for that KC (*9, 44*). The simulations revealed that α/β KCs excelled in detecting class 3 odors, while γ KCs demonstrated superior sensitivity to class 1 odors (Fig. 4A). For the assessment of coding capacity, we examined the KC hamming code dimensionality, using binary representations to describe activated and inactivated states (*45-47*). Notably, through PCA, we found that γ KCs exhibited significantly higher coding capacity compared to α/β and α’/β’ KCs (Fig. 4B). In summary, the complex hybrid network architecture offers a range of advantages: α/β KCs exhibit heightened sensitivity to class 3 odors, while γ KCs excel in detecting class 1 odors. This architecture also maintains high coding capacity, potentially enabling refined odor discrimination.

**Fig. 4.**
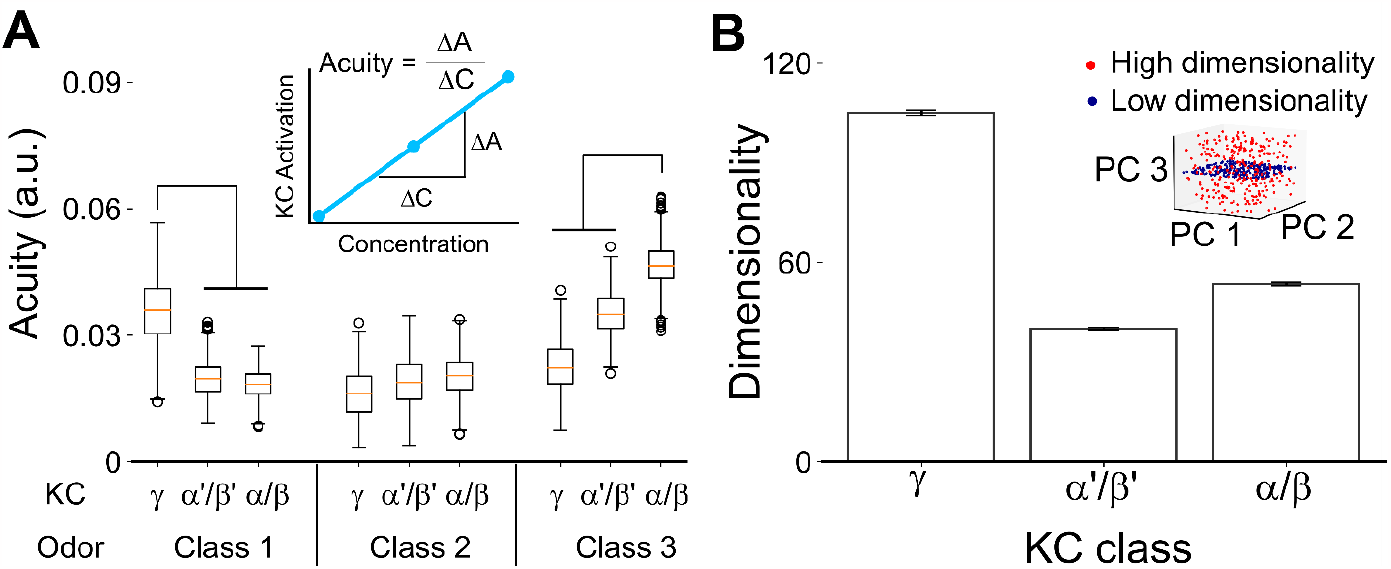
Hybrid network architecture diversifies olfactory coding strategies. (**A**) Olfactory acuity among KC classes to artificially generated odors in three odor classes by numerical simulation (see Methods). Friedman test with Dunn’s post hoc test was used for pairwise group comparisons. For each comparison, the p-value was < 0.001. The inset illustrates the calculation of odor acuity, which is the change in KC activation ratio over different odor concentrations. (**B**) The coding capacity among KC classes for artificially generated odors is defined by dimensionality (see Methods). One-way ANOVA with Tukey’s HSD was used for pairwise group comparisons. For each comparison, the p-value was < 0.001. The inset shows a simplified depiction of coding capacity measured by dimensionality.

## Discussion

The MB serves as a pivotal hub in processing a vast array of olfactory cues, with specific emphasis on critical food and reproductive scents. Our detailed analysis of the connections between PNs and KCs within the MB’s calyx has illuminated an intricate hybrid architectural design that seamlessly accommodates these functional roles. Particularly noteworthy is the selective interaction pattern observed between γ KCs and cluster 1 PNs, integral for pheromone sensing, while α/β KCs establish preferential links with cluster 3 PNs associated with food-related odors. This precise wiring specificity emerges from the spatial convergence of PN boutons and KC claws, strategically positioned within specific regions of the calyx as suggested previously (*17, 18*). It is worth highlighting that this distinct connectivity motif observed between cluster 1 and cluster 3 PNs is consistently maintained across the AL, calyx, and LH. The informative simulations, driven by these established connections, provide valuable insights into the diverse responses of KCs to various odors. These simulations notably emphasize the shared sensitivities exhibited by cluster 3 PNs. Beyond this, the simulations underscore two discernible strategies: γ KCs adopting a more stochastic wiring pattern, promoting the capacity of odor coding, while α/β KCs leverage a stereotypical architecture to enhance detection sensitivity, particularly attuned to food odors. Furthermore, our investigation also reveals an intermediate profile in α’/β’ KCs, with α’/β’-m resembling α/β KCs, favoring short-chain esters, while α’/β’-ap seems displaying a preference for aroms. In essence, our study provides a remarkable glimpse into the intricate hybrid wiring strategy harnessed by the *Drosophila* MB. This strategy effectively marries acuity and capacity, enabling the MB to adeptly process a diverse array of olfactory cues, including those pivotal for survival and reproduction.

Comparing the realistic connections with their shuffled counterparts has yielded a rich understanding of the diverse input preferences that govern the connections between distinct KC classes and their corresponding PN clusters. The meticulous shuffling of connection targets while maintaining consistent PN-to-KC connection numbers has brought to light intriguing connection ratios, distinctly originating from PN cluster 1 and cluster 3 toward downstream KCs. This contrasts with the somewhat obscured preferences noted in the examination of stochastic shuffling, as previously studied (*22*). Notably, recent investigations employing patch-clamp techniques and dye-filling methods have seemingly inclined toward advocating for a more adaptive, experience-driven “random” wiring model (*20*). These disparities with earlier findings might stem from potential sampling biases, especially toward γ KCs, owing to their comprehensive inputs encompassing all AL glomeruli. Through our comprehensive simulations, we have underscored the significance of having a substantial representation of KCs – at least 100 KCs for a given KC class with complete connection data – in order to unveil the broader global input preferences of that particular KC class.

Transitioning to the functional aspect, earlier research hinted at a lack of stereotyped responsive profiles within α/β-c KCs across different individual animals at the single cell level (*24*). However, functional results show predictable tuning features of KCs in both axonal bundle regions *(11)* and soma areas (*25*). Our model prediction followed by calcium imaging validation uncover the preferential spatial connectivity at population level. Therefore, even the stereotyped tuning profile is absent for individual KCs, the soma distribution (*25*), tuning profiles of KCs at the lobe level, and potentially the mushroom body output neuron (*11*) can still maintain the preferences across individuals.

Moreover, even though a random connection model can generate conserved clustered odor representations (*23*) and stereotyped readout (*48*) by the mushroom body output neurons, the divergent tuning profiles of different KC classes to aromatic versus aliphatic odors at population level rely on, at least, class-based connection preference. Our study not only solidifies the notion of odor response stereotypy but also serves to validate predictions pertaining to specific odors, exemplified by odors like EP and PA that induce notably stronger responses in α/β KCs compared to their γ counterparts – a trend consistently affirmed through calcium imaging across multiple adult flies. This line of evidence leads us to suggest that the connection preferences discerned from the FlyEM dataset (*31*) extracted from a single fly likely extend beyond the individual and encapsulate broader patterns within the fly population.

The functionality of both artificial and biological neural networks hinges upon their underlying network architectures, playing pivotal roles in tasks ranging from discrimination to generalization (*15, 49*). Notably, recent computational endeavors have drawn intriguing parallels between the intricate circuitry of the MB and the concept of hashing – a mechanism that transforms the multidimensional chemical space into sparse codes, thereby facilitating processes like novelty detection and generalization (*47, 50*). In the realm of neural networks, the role of randomness in shaping connectivity is critical, as random networks tend to offer heightened coding dimensionality (*51*), thereby potentially optimizing the discrimination capacity of the system.

In light of our investigation, a novel architecture emerges – the hybrid MB network – which exhibits distinct patterns of connectivity among different KC classes. Specifically, the α/β KCs showcase a preference for integrating inputs from cluster 3 glomeruli, fostering the repetition of specific input combinations. On the other hand, the γ KCs emerge as universalists, receiving inputs more uniformly from all glomeruli, thus positioning them as adept discriminators. This unique wiring configuration renders γ KCs remarkably proficient in odor discrimination, attributed to the inherently stochastic nature of their connections.

The preferential connectivity between PNs from cluster 1 glomeruli and γ KCs sheds light on the critical role played by these connections in shaping behaviors related to courtship (*52, 53*). Conversely, the α/β KCs, by virtue of their convergent input, might sacrifice some resolution, yet the presence of specific connections significantly bolsters their ability to detect food-related odors. Furthermore, the clustered responses enhance the ability for odor generalization (*23*). This intricate hybrid network architecture can be construed as a more refined model, finely attuned to satisfy the fly’s complex demands in terms of sensory sensitivity and discrimination, particularly when it comes to processes like learning and memory. In essence, our findings underscore the exquisite balance between sensitivity and discrimination achieved through the hybrid network configuration, offering insights into how neural architecture can be optimized to meet distinct functional demands.

While we focused on the three major KC classes and several subclasses, they could be further subdivided into more families that play distinct roles in olfactory learning and memory processes (*10, 54-58*). The exact PN connectivity preferences specific to these KC families remain unexplored. The role of APL neurons innervated the entire MB in suppressing KC sparse coding using gamma-aminobutyric acid is not fully understood and whether this suppression follows a random pattern is uncertain (*7, 59*). Additionally, our analysis did not delve into KCs’ connectivity preferences with multiglomerular PNs, crucial for cross-glomerular integration (*60*). Moreover, it remains unclear whether sister PNs originating from the same glomerulus contribute to the diverse sparse coding patterns of KCs, as observed in the calycal spatial map of pheromone-sensing PNs in cockroaches (*61, 62*). Addressing these aspects would provide a more comprehensive understanding of the intricate information processing within the MB circuitry.

## Materials and Methods

### Data sampling

This research is based on the FlyEM dataset v. 1.2.1 (*31, 63*), a fruit fly’s hemibrain connectome with image resolution down to the synaptic level. The connection between a PN and a KC was determined by at least three synapses. Since we focused on investigating the olfactory encoding mechanism of *Drosophila melanogaster*, only 52 types of olfaction-related glomerular PNs were selected. We excluded PNs from VP-type glomeruli, multiglomerular PNs and KCs without receiving any connection from the monoglomerular PNs from the 52 glomeruli. Eventually, we selected 106 PNs and 1745 KCs on the right side of the brain from the FlyEM dataset (*31*). KCs can be categorized into three major classes (*54*) – γ, α’/β’, and α/β or subdivided into subclasses (*31*) including γ-m, γ-super, γ-d, α’/β’-ap1, α’/β’-ap2, α’/β’-m, α/β-m, α/β-c, α/β-p, and α/β-s. Considering that the synapses of the PN-to-KC network demonstrate high plasticity and may differ among individuals (*29*), we defined the connection between a single PN and KC as the connectivity unit (neuronal level). The selected PN-to-KC connectivity is shown in (Fig. 1a).

### Connection preference evaluation

The preference score *G*_*k*_ for connections from PNs of a glomerulus *g* to a KC *k* is defined as,

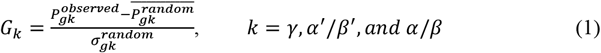

where 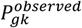 is the ratio (0-1) of the connection from a given glomerulus *g* to the KC class *k*. 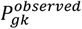 is normalized such that 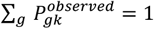 for each *k*. 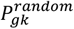 measures the same quantity but for the randomized glomerulus-to-KC network. Specifically, the targeted KCs are randomly chosen for every glomerulus, but the number of targeted KCs remains the same. The process is repeated 1000 times to calculate the mean innervating ratio 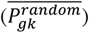 and the standard deviation 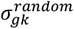. *G*_*k*_ within the interval of ± 2 is considered as no significantly preferred projection from the glomerulus *g* to the class-k KCs.

### Global shuffling for connections

We compared the observed data (FlyEM dataset (*31*)) with 1000 random datasets generated by the shuffling algorithm. The shuffled ratio, *x%*, is given from 0% to 100%. For the full shuffling process (*x* = 100), all glomerular PNs were reassigned with new downstream KCs at random. The effect of the full shuffling process was identical to the shuffling algorithm described in Caron et al., 2013 (*20*) maintaining the total connection numbers of each glomerulus and each KC. For partial shuffling, only *x%* of connections in the FlyEM dataset were randomly selected and shuffled.

### Local shuffling for connections

Compared to the global shuffling, we further set spatial constraints. After identifying the bouton/claw structure by DBSCAN, we measured the distance between every two claws, and exchanged the upstream boutons if the distance of the claws were shorter than *R*. The same process was repeated for 1000 times. The total numbers of connections, boutons, and claws were preserved under local shuffling. For the schematic plot, please see fig. S15.

### Connectivity correlation quantification

We applied Pearson correlation analysis to measure the downstream connection similarity of each two glomeruli. The connection profile of a single glomeruli is regarded as a vector which is mapped into a 1735-dimensional space (the total number of KCs). The Pearson correlation is calculated by

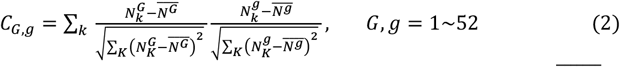

where 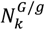 is the total connected PN number between *G*^*th*^*/g*^*th*^ glomerulus and *k*^*th*^ *KC*, and 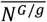 is the average downstream connection number of *G*^*th*^*/g*^*th*^ glomerulus. To visualize the convergent glomerular clusters, we implemented *Ward* hierarchical clustering using Python’s Scikit-Learn library.

### Input preference quantification

To estimate the degree of input preference of each KC class, the correlation based principal component analysis (PCA) was implemented by calculating the eigen values of the Pearson correlation matrix of the glomerulus-to-KC connection matrix using Python’s Numpy library. The set of principal components (PCs) is a linear transformation of the connection matrix that generates maximum variance, and the percent variance of the first PC (PC1) is correlated with the diversity of the input combination. If more KCs receive similar glomerular input combinations than the random hypothesis suggests, the explained variance ratio of the PC1 of the glomerulus-to-KC network would be larger than that of a completely shuffled network. We independently shuffled the connections within each of the three KC classes, varying the shuffled ratio from 0% to 100%. The degree of preference is defined as the minimal shuffled ratio *x%* that satisfies

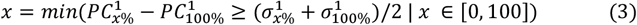

where 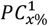 and 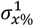 are the mean and standard deviation of the explained variance of PC1 calculated from 1000 partial shuffled connectivity trials.

### Categorical shuffling for neuronal classification

We utilized a categorical shuffling algorithm to explore whether different classes of KCs and clusters of PNs occupy distinct regions within the calyx. This algorithm allowed us to compare the actual distribution of KCs and PNs with a distribution derived from a null model. The null model was created by randomly shuffling the classification of neurons (the class of a KC or the cluster of a PN) while maintaining the total number of neurons in each category. For example, when examining KC classes, such as γ, α’/β’, and α/β, we randomly reassigned the KC class for each KC while keeping the overall number of neurons in each class constant.

### Categorical shuffling for claw/bouton

For claw/bouton analysis, we built a null model by randomly shuffling each neuron’s claw/bouton distribution while preserving the total number of claws/boutons associated with each neuron. For instance, consider a KC endowed with five claws. Upon implementing the shuffling algorithm, this KC will still possess five claws, but it will exchange them randomly with other KCs, resulting in a different arrangement.

### Categorical shuffling for synapse

For the synaptic site analysis, we extended our methodology by implementing a null model specifically designed to investigate the distribution of synapses between neurons. To create this null model, we employed a random shuffling algorithm that preserved the total number of synaptic sites associated with each neuron. In particular, we focused on the hypothetical scenario where neuron *A* establishes connections with neuron *B* at only one specific site. To account for this, we treated all synapses between neurons *A* and *B* as a single unit during the permutation process. By doing so, we ensured that the overall number of synaptic sites remained consistent while introducing random shuffling.

This approach mirrored the algorithm utilized for the claw/bouton analysis, where we maintained the integrity of the total number of synaptic sites per neuron while shuffling their distribution.

### Odor selection

We collected the response profiles of 52 olfaction-related odor receptors (ORs) from the DoOR database (*42*). Each glomerulus connects with a distinct type of olfactory receptor neurons (ORNs), and most ORNs only express a single type of OR (*64*). The DoOR dataset includes the OR’s response to 693 odors, which is normalized by the following equation:

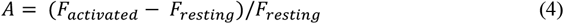

where *A* is the response activity. *F*_*resting*_ is derived from the response to mineral oil. *F*_*activated*_ is derived from the response to other odors. The missing values in DoOR are manually converted to “0” in our study.

### Simulation for odor response in MB lobe

The odor response of the MB lobe is closely correlated with the activity of its upstream glomeruli and the strength of their connections. Each KC class has a distinct average claw number, and KCs with a higher claw number demonstrate a higher activation threshold (*9*). As a result, the KC response level has a higher correlation with the percentage of claws than the absolute number of claws that receive inputs. Therefore, we assume that the KC response level is associated with the activated ratio of KCs, denoted as *R*_*gk*_ and given by

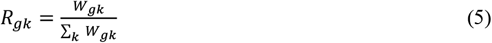

where *W*_*gk*_ represents the number of connected claws between glomerulus, g, and a single KC, k. *R*_*gk*_ is a normalized value, which is divided by the total claw number of each KC.

The simulated odor response of the MB lobe, represented as 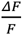, is determined by the summation of inputs:

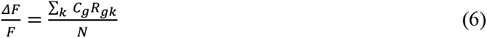

where *N* represents the total number of KCs in a specific class, and *C*_*g*_ is the activity of the upstream glomerulus, *g*. Based on the one-to-one mapping structure, we assume that the glomerular activity is proportional to the activity of its upstream ORNs, and the level of ORN odor-evoked activity refers to the data from the DoOR database (*42*).

### Spatial distribution similarity analysis

To compute distribution similarity, the first step was to collect all the skeleton points, center points of boutons/claws, and synapse coordinates according to their KC classes or PN clusters. Then, we attained the spatial distribution by kernel density estimation, KDE, using Python’s Scipy library. Next, we calculated the Jensen–Shannon divergence, JSD, to estimate the distribution similarity using the SciPy package as well (*65*). JSD’s advantages include (1) boundedness and (2) symmetry. Therefore, it helped us normalize the similarity among different distributions. JSD is defined by

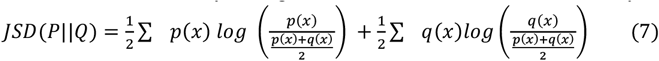

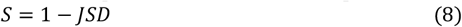

where *P* and *Q* are the distribution of *p* and *q. S* is distribution similarity. The spatial distribution preference was evaluated by the difference in distribution similarity between the original data and the shuffled ones. Categorical shuffling was achieved by switching neuronal classification between different individual neurons. For synapses, we hypothesized that neuron A connects neuron B at only one site. Thus, all synapses between A and B were packed as one unit to do permutation.

### Bouton/claw identification

To identify the middle-scale anatomic structures – boutons for a PN and claws for a KC – we used DBSCAN from Python’s SciPy library. The minimum number of points was set to 3, which is consistent with our connection threshold, and the radius was set to 1.6 µm. Since one bouton connecting to several KC claws should not be identified as several boutons. After manual inspection, we established a distance threshold of 2.8 µm. If the distance between the centers of two boutons/claws from a PN/KC was less than 2.8 µm, these two structures merged into a single entity automatically. The example results are presented in fig. S1B.

### Weight ratio calculation

The input weight ratio was calculated based on the connection weight from a PN cluster to a KC, divided by the total connection weight for the KC from all PNs. Specifically, the input weight ratio is given by the formula:

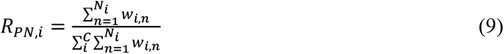

where *R*_*PN,i*_ is the input weight ratio of a KC class receiving from a cluster *i, N*_*i*_ is the total number of PNs in cluster *i, C* is the number of clusters, and *w*_*i,n*_ is the connection weight of a PN to a KC.

As for the output weight ratio, it was calculated using the connection weight for a PN to a KC class, divided by the total connection weight for a PN to all KC classes. The value is defined as follows:

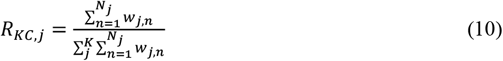

where *R*_*KC,j*_ is the output weight ratio of a PN cluster to a KC class *j, N*_*j*_ is the total number of KCs in KC class *j, K* is the number of KC classes, *w*_*j,n*_ is the connection weight of a PN to a KC.

### Correlation of weight ratio and synaptic distribution preference

To evaluate the correlation between the connection weight ratio and the spatial distribution, we calculated the differences in the connection weight ratio (for each KC class to each PN cluster) between the original network and the shuffled connection ratio. Next, we calculated the differences in the spatial distribution preference by subtracting the shuffled distribution similarity from the original distribution similarity. Finally, the correlation between the difference in the connection weight ratio and the difference in distribution similarity was estimated by performing linear regression using Python’s SciPy package.

### Weight normalization for simulation

Previous studies show that the KC activation is correlated with the percentage of the claws that receive inputs (*9, 44*). Therefore, we normalized the connection matrix W_PN>KC_ by the column. As a result, each connection weight between a PN to a KC is given by

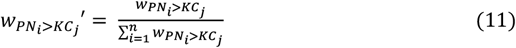

where *n* is the total number of PN and w_PNi>KCj_ represents the original connection weight (synapse number) of PN *i* to KC *j*.

### Simulation for artificial odors

To investigate how the connection preference affects KC coding, we generated artificial odors which only activate glomeruli within the same glomerular cluster. Here, we randomly choose *m* candidate glomeruli from *n* glomeruli and define the activity *A*_*PN*_ of the PNs from the chosen glomeruli as

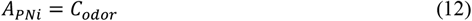

Otherwise,

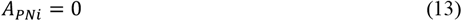

where *C*_*odor*_ is set from 7 to 11 to observe how odor stimulation strength affects KC activation. The unit of *C*_*odor*_ is arbitrary. Note that the absolute value of *C*_*odor*_ is not important as it can always be compensated by the scaling factor and activation threshold introduced below. The range of *C*_*odor*_ is used to simulate the firing rate dependency of PNs to the odor concentration. For odors that only activate cluster 1 glomeruli, we defined the odors as class 1 odors. For each odor class, we generated 1000 different odors without the same activated glomerular combination.

### Simulation for KC representation of artificial odors

To investigate how the connection preference affects KC coding, we used the artificially generated odors with the observed connection matrix derived from FlyEM dataset (*31*) to analyze the KC activation profile. The activity of KC is given by

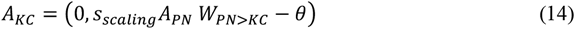

where *θ* represents the activity threshold, which was set to 1. *A*_*KC*_ is the vector representing KC activity for each odor filtered through the ReLU function. *A*_*PN*_ is derived from the artificial odors mentioned above. *s*_*scaling*_ is a scaling factor that was set to 0.3. *W*_*PN>KC*_ corresponds to the connection matrix. Here, we carefully adjusted *s*_*scaling*_ and *θ* to achieve a KC activation ratio of approximately 10% when the *C*_*odor*_ is 8. This ratio is in line with previous experimental studies (*43*).

### Acuity analysis

To evaluate olfactory acuity for different odor classes, we first assessed the activation ratio of KCs. The activation ratio was calculated by dividing the number of activated KCs by the total number of KCs in response to each odor stimulus.

Next, we analyzed the relationship between the odor stimulation strength (*C*_*odor*_) and the activation ratio by performing linear regression using Python’s Scipy library. This regression analysis allows us to estimate the slope, which represents the change in the activation ratio per unit change in odor stimulation strength (*C*_*odor*_). The odor acuity for a KC class to an odor is defined by the slope of the regression line.

### Dimensionality analysis

To compute the dimensionality of KC representation, we performed PCA decomposition for the KC response matrix. The dimensionality is defined by (*51*)

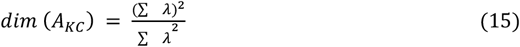

where *λ* is the eigen value. For hamming code capacity calculation, all positive A_KC_ values were set to 1.

### Fly strains

All fly stocks were reared on standard corn meal-yeast-agar medium at 25 °C or 18 °C with around 70% humidity under a 12:12-h light/dark cycle. The following fly lines were used in the current study: *OK107-Gal4* drives the expression of all three classes of MB neurons, *VT57244-Gal4* (v200970, Vienna Drosophila Resource Center) for the α’β’ neurons, *UAS-GCaMP7f* (79031, Bloomington Drosophila Stock Center) flies carrying a transgene for a genetically encoded calcium sensor.

### Chemicals and stimulation

For olfactory stimulation, ethyl propionate (EP, CAS: 105-37-3, Sigma-Aldrich), isopentyl acetate (IA, CAS: 123-92-2, Sigma-Aldrich), octyl acetate (OA, CAS: 112-14-1, Sigma-Aldrich), pentyl acetate (PA, CAS: 628-63-7, Sigma-Aldrich), benzaldehyde (BA, CAS: 100-52-7, Scharlau), anisole (ANS, CAS: 100-66-3, Scharlau), pyrrolidine (PYR, CAS: 123-75-1, Sigma-Aldrich), and ethanol (EtOH, CAS: 64-17-5, Honeywell) were used. Odors were diluted in mineral oil. The concentration was 10^−6^ (v/v) for a single concentration test. Concentration gradient experiments were adjusted from 10^−6^ to 10^−2^ (v/v). The diluted solution was packed in glass tubes at a volume of 10 ml per tube. Using a computer to control the olfactometer, the odor was introduced into a continuous flow of air for the antennae to detect. Stimulus duration was 10 sec. Airstream continued for the entire experiment time. Flies were stimulated with different functional group odorants (ester, non-ester, and arom) and two blank controls (mineral oil and ethanol).

### Confocal imaging and imaging processing

The sample brains were imaged by Zeiss LSM 710 or LSM 780 confocal microscopy with a 40× C-Apochromat water immersion lens. Images were scanned under the following setting: 512 × 512-pixel resolution, scanning speed 7, and line average 2 in ZEN software (Zeiss). Max-projection images and single-plane images were formatted using ZEN software (Zeiss).

### In vivo GCaMP functional imaging and imaging processing

Flies expressing *UAS-GCaMP6* in α’/β’ KC and *UAS-GCaMP7* in whole KC were mounted on a droplet-shaped sheet, with a window opened on the head capsule, and then adult hemolymph-like (AHL) saline (108 mM NaCl, 5 mM KCl, 2 mM CaCl2, 8.2 mM MgCl2, 4 mM NaHCO3, 1 mM NaH2PO4, 5 mM trehalose, 10 mM sucrose and 5 mM HEPES, pH 7.5, 265 mOsm/kg H2O) added immediately.

Recording of changes in GCaMP intensity before and after odor stimulations was performed on Zeiss LSM 780 with a 40× C-Apochromat water immersion lens. Images were acquired with 512 × 512 pixel resolution at two frames per second for 60 frames. For odor stimulation, odorants were delivered during 10 – 20 sec for odor stimulation in each 30 sec trial. Each odor was presented with an inter-stimulus interval of 1 min.

A 488 nm LED light source was placed beneath the chamber to provide red-light stimulations. MB lobes were imaged in an area of the dorsal surface of the fly brain. The ROIs of α/β KC and γ KC were circled at the tip of the lobe, while α’/β’-m KC and α’/β’-ap KC were circled according to the position of the nerve distribution provided by neuPrint (*63*).

### Data analysis for functional imaging

The raw fluorescence signals were converted to *ΔF/F*, where *F* is the averaged baseline fluorescence value of 10 sec before the stimulation onset, and *ΔF* is the difference between the highest signal and the mean. Peaks of post-response were taken as maximum *ΔF/F* within the 10 sec following light onset, compared using the nonparametric Wilcoxon Signed Rank Test.

### Statistics

Statistical analysis was performed by Microsoft Excel®, GraphPad Prism®, and Python. Regression analysis was used to capture the correlation between two variables. The non-parametric Wilcoxon Signed Rank Test was performed to compare the calcium responses between two lobes. For non-normal distributed group data, Friedman test with Dunn’s post hoc test was used. Otherwise, we performed one-way analysis of variance (ANOVA), with Tukey’s HSD. The significance thresholds are indicated by *: p < 0.05, **: p < 0.01, and ***: p < 0.001. All data are presented by mean values ± SD.

## Acknowledgments

We would also like to express our thanks to the Bloomington Drosophila Stock Center and the Vienna Drosophila RNAi Center for generously providing the fly stocks used in our research. We extend our gratitude to Li-An Chu for providing valuable insights and feedback on our study.

## Author contributions

Conceptualization and Supervision: ASC, TKL, CCL, KLF. Methodology and Investigation: LSC, CCC, RHC. Software, Resources, Formal analysis, and Visualization: LSC, CCC. Writing—original draft: LSC, CCC. Writing—review & editing: KLF, ASC, CCC, CCL, LSC, TKL.

## Data and materials availability

All data and materials are available in the main text or the supplementary materials. Codes are available upon the request made to the first authors or the corresponding authors.

**Fig. S1.**
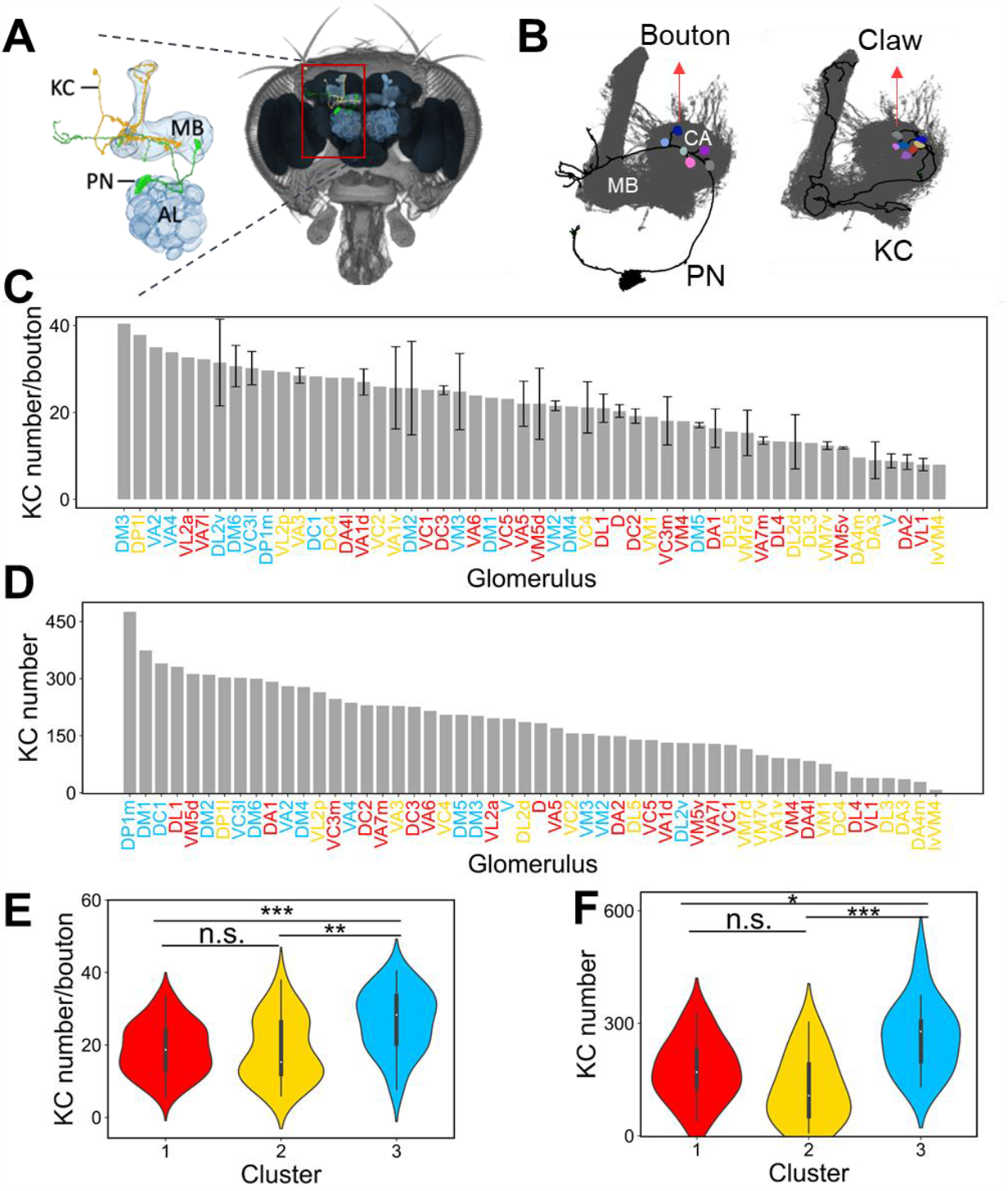
Differential bouton capacity of glomeruli for KCs. (**A**) A *Drosophila* brain (right) and a zoom-in view of the AL and MB (left). A pair of representative PN (green) and KC (yellow) makes synaptic contacts in CA. (**B**) Boutons from one representative PN (left) and claws from one representative KC (right). (**C**) The average number of downstream KCs connected by each bouton from a given glomerulus. The color of each glomerulus is assigned by cellular connection preferences in Fig. 1B. Each error bar represents the standard deviation across individual PNs from the given glomerulus. (**D**) The total downstream KCs connected by each glomerulus. The color code is the same as in (**C**). (**E**) The distribution of KC number/bouton and (**F**) total connected KC number of glomeruli in PN clusters. (Kruskal-Wallis one-way ANOVA with Bonferroni’s post hoc test, and significance levels were indicated as follows: n.s. for not significant, * for p < 0.05, ** for p < 0.01, and *** for p < 0.001.).

**Fig. S2.**
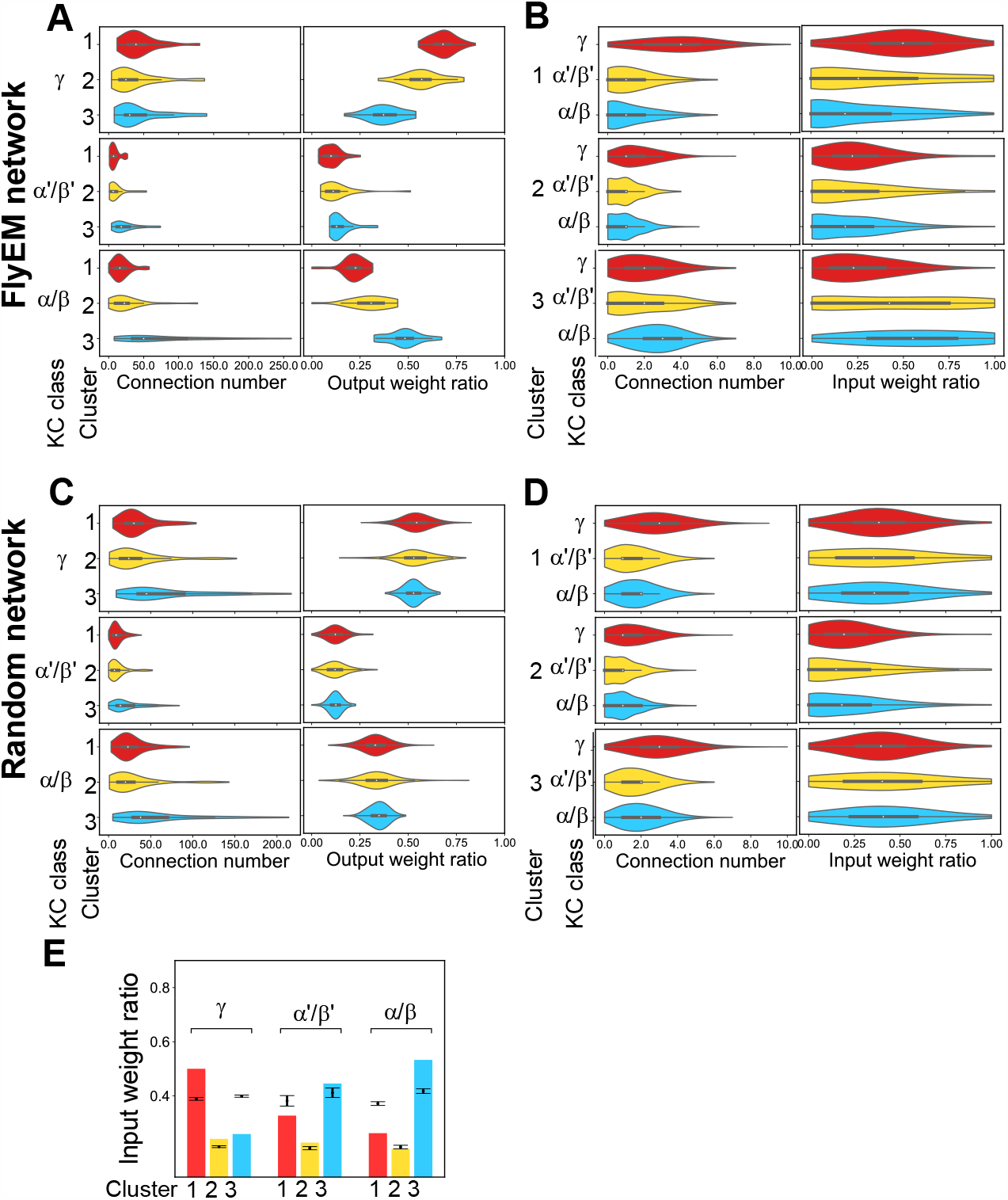
Summary of PN-to-KC connections. (**A**) The distributions of PN connection number (left) and output weight ratio (right) to each KC class, γ (top), α’/β’ (middle), and α/β (bottom) for PNs in each cluster. The output weight ratio is defined as the weight of a PN to a KC class divided by the total weight of the PN to all KCs (see Methods). Note that the PN cluster 3 forms significantly more connections to α/β KCs than other PN clusters. (**B**) Similar to (**A)** but from the KC perspective. The connection numbers represent the number of PNs (in a cluster) from which a KC receives input. The input weight ratio is defined as the weight of a PN cluster to a KC divided by the total weight for a KC from all PNs (see Methods). Note that γ KCs receive significantly more connections from the PN cluster 1 than other KC classes. Part of the data is displayed in Fig. 1C, and the full data is shown here. (**C**) and (**D**), same as (**A**) and (**B**), respectively, but for shuffled networks. The outgoing connection number for each PN and the incoming connection number for each KC are preserved during shuffling. (**E**) The input weight ratio of three KCs from PN clusters. The error bar indicates the result from 30 trials of shuffling.

**Fig. S3.**
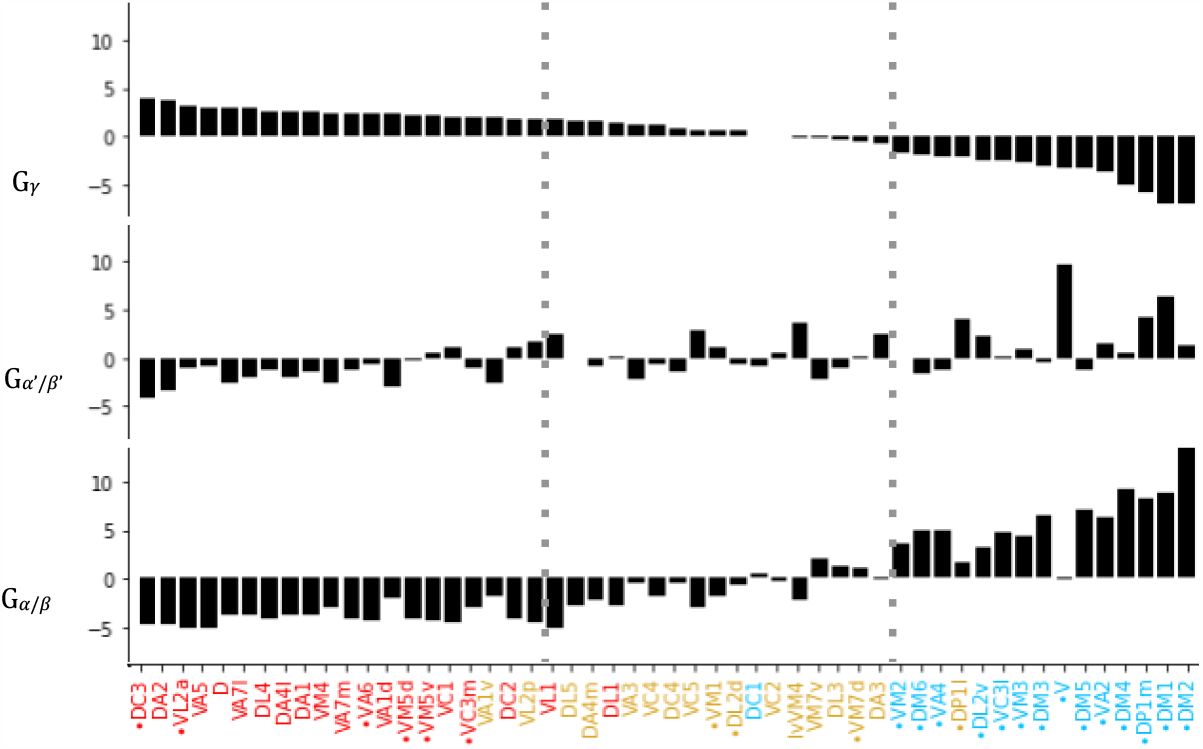
The preference score G_i_ for each glomerulus with the synaptic numbers taken into consideration. Instead of counting each connection between a PN and a KC as one (as in Fig 1), in this plot, the connection between a pair of PN and KC is weighted by the number of synapses formed between them. Three preference clusters are color labeled with red (cluster 1), yellow (cluster 2), and blue (cluster 3).

**Fig. S4.**
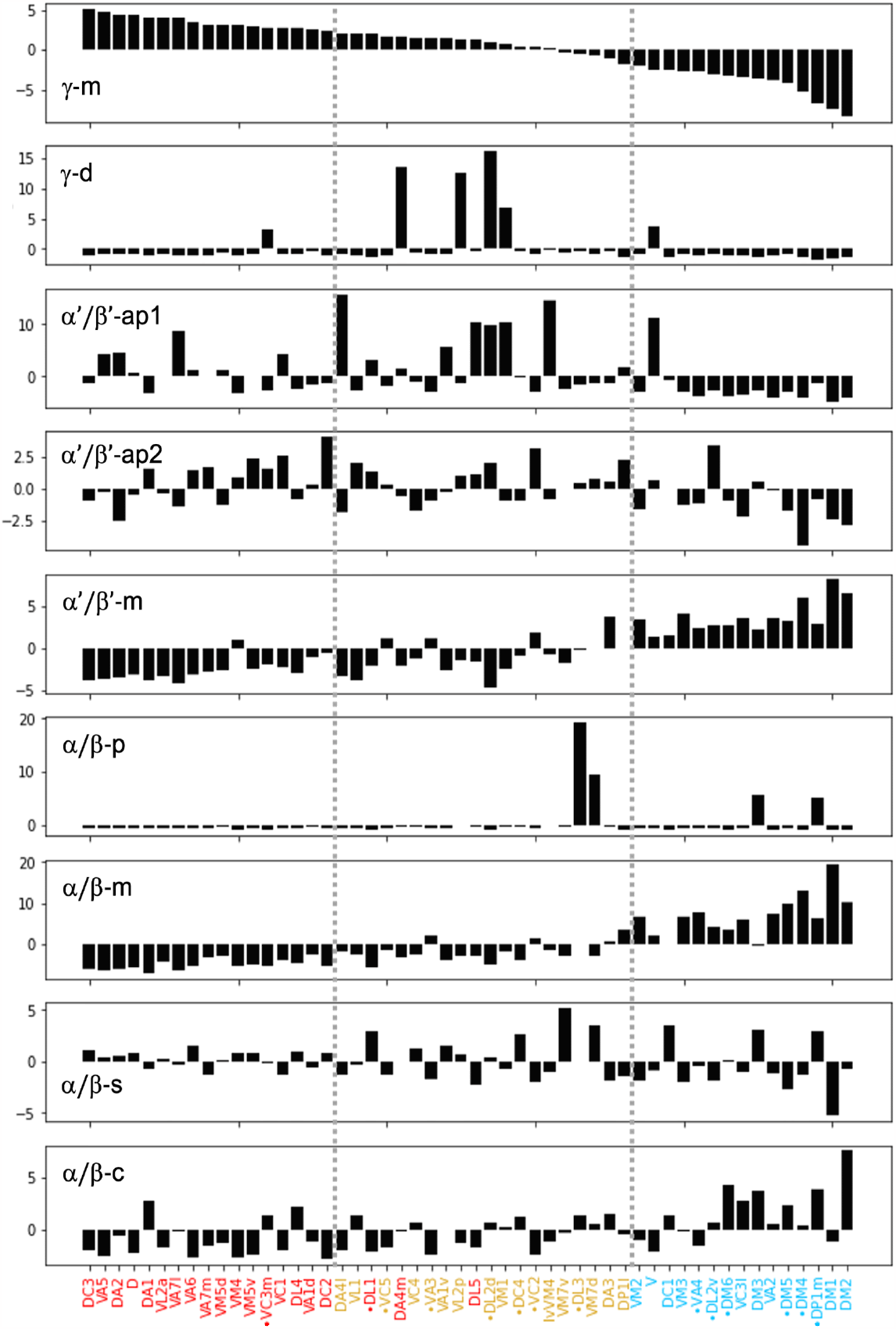
The preference score of KC subclasses. The glomeruli are sorted by the preference score values of γ-m. Three preference clusters are color labeled in red (cluster 1), yellow (cluster 2), and blue (cluster 3).

**Fig. S5.**
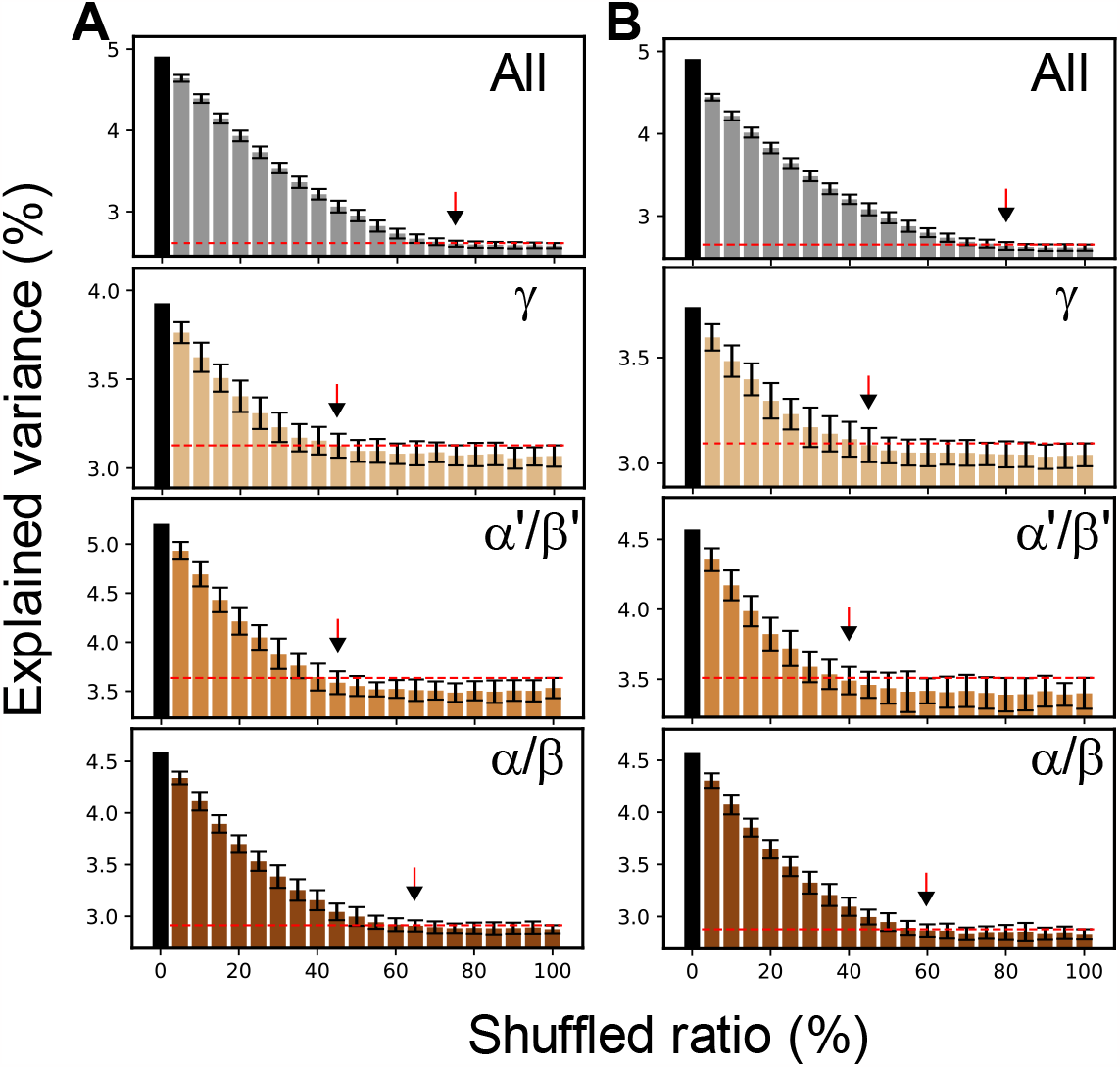
The non-random structure in the PN-KC network as revealed by random shuffling. (**A**) The explained variance of the first principal component of the PN-KC networks containing all KCs, γ, α’/β’, or α/β KCs (from top to bottom) with partially shuffled connections. The result implies a non-random structure in the PN-KC network. The error bars indicate the standard deviation of the explained variance from 1000 shuffling trials. The red dashed line indicates the standard deviation under 100% shuffling. Arrows mark the shuffled ratios that are required to remove the non-random structure. (**B**) Same analysis as in (**A**), respectively, but the connections are weighted by the synaptic numbers between PNs and KCs.

**Fig. S6.**
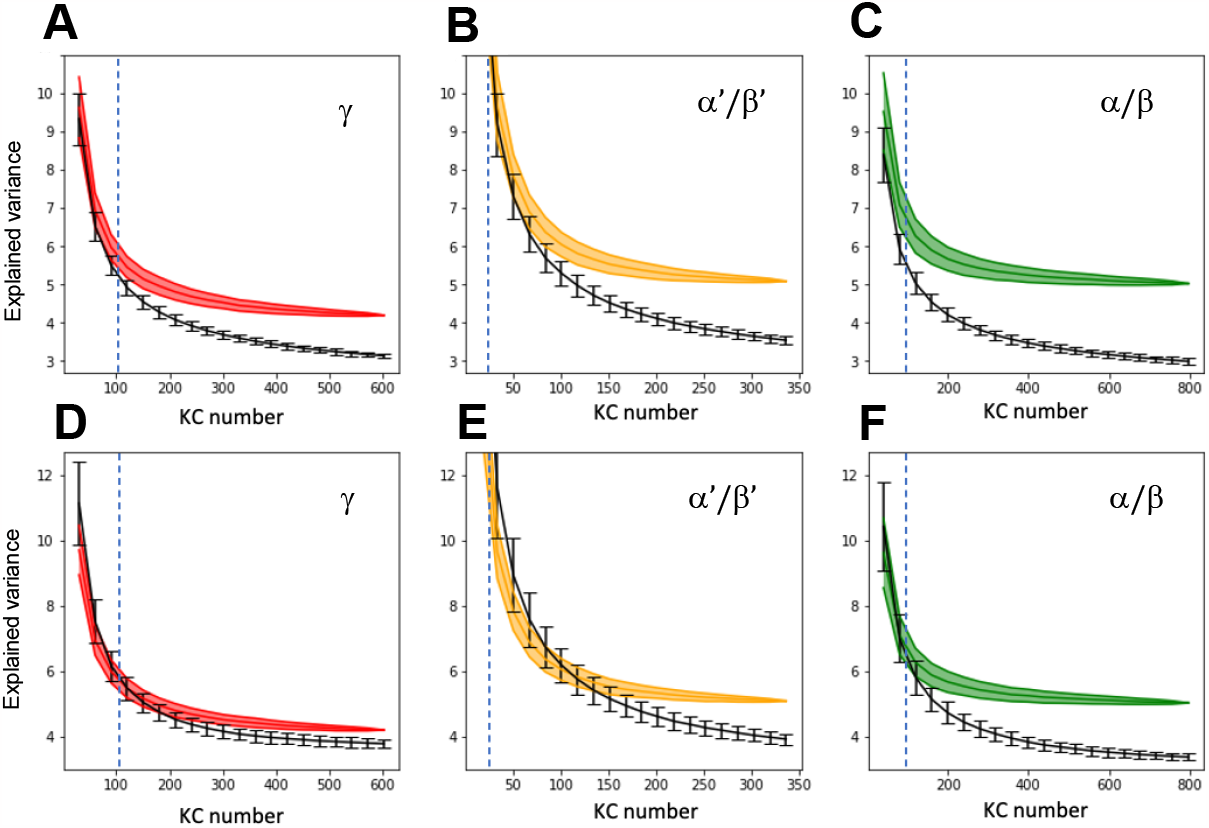
Partial sampling of the KCs is less likely to reveal the non-random structure in the PN-to-KC network. (**A**-**C**) Comparison of the explained variance of the first principal component (y-axis) between the PN-to-KC network (colored curves) and the shuffled network (black curves) under the same degree of partial sampling (x-axis). The shuffling is performed with the preservation of the total KC number and the connection number of each KC and each glomerulus. The color-filled region represents the mean value ± s.d. of the explained variance from 1000 sampling trials. The claw number of each KC in observed data is also preserved. The error bar represents the mean value ± s.d. of the percent variance from 1000 shuffling trials. The blue dashed lines mark the sampling sizes in Caron et al., 2013 (*20*). (**D**-**F**), The same procedure is applied on the partially sampled PN-to-KC network and shuffled networks, but with 50% of the connections randomly excluded. The result shows that a large sampling size is required to discern the difference between a partially sampled network and shuffled networks.

**Fig. S7.**
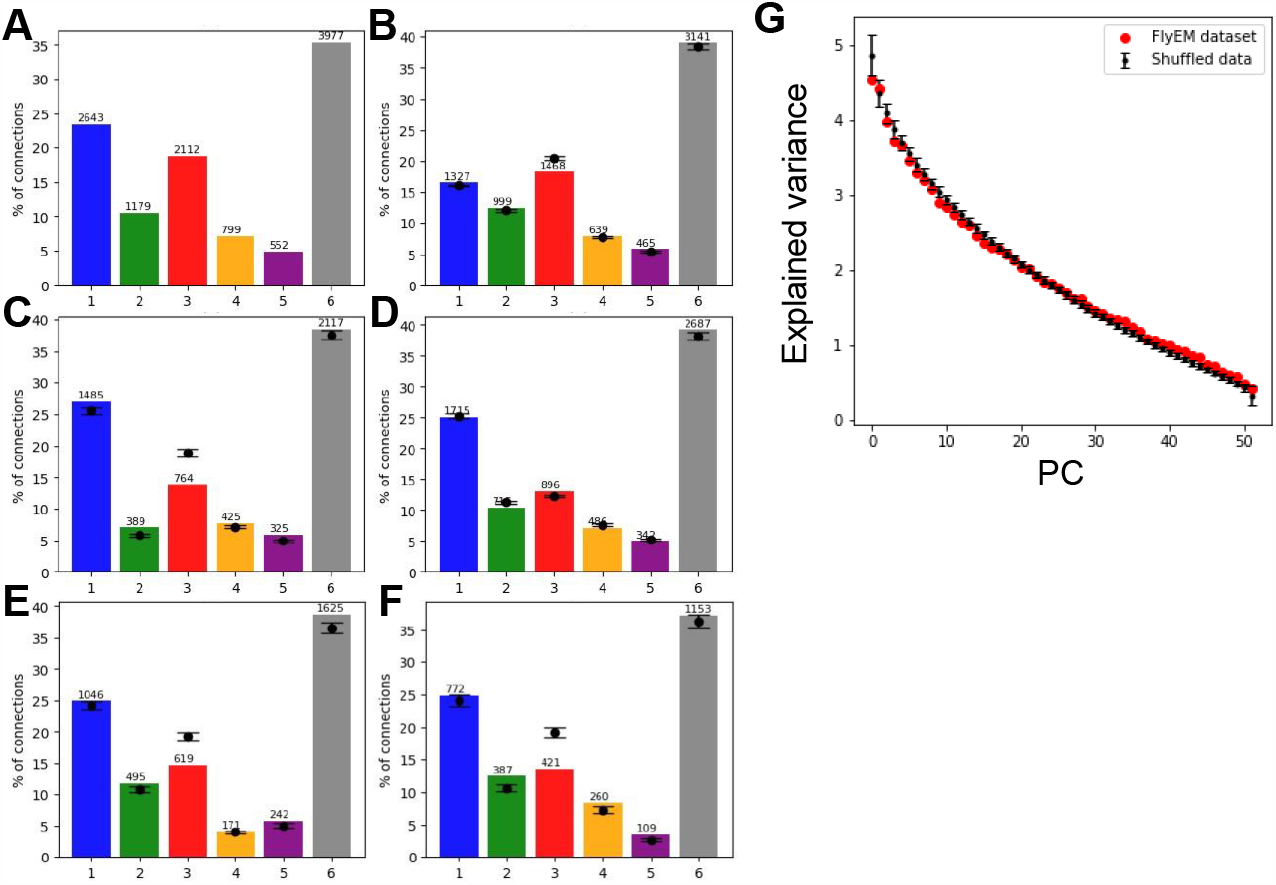
Random-like network structure due to incomplete sampling of the PN-to-KC network. We conducted the group analysis of PN-to-KC connections following the same procedure performed in Caron et al. (2013) (*20*). The glomeruli are classified into five groups based on their PNs’ anatomy in the lateral horn (*18*) (Table S1). The remaining glomeruli are classified as the group ‘other’ (grey column). (**A**) The percentage of PN connections (to all KCs) from different glomerular groups. The number of connections is labeled above the bars. (**B**) Similar to (**A**) but for the PN connections to the subset of KCs that receive a least one connection from glomerular group 1. The same analysis is also done with shuffled networks and is indicated by the dots (mean) and the error bars (standard deviation). (**C**-**F**) Same as (**B**) but for PN connections to subsets of KCs with at least one connection from glomerular group 2∼5, respectively. (**G**) The variance explained by each principal component, PC, in PCA for the partially sampled PN-to-KC network with 50% of input dropped out and subsets of KCs (102 γ KCs, 14 α’/β’ KCs, and 84 α/β KCs) selected. The variance distribution of the subsampled connections is indiscriminate from the shuffled connections.

**Fig. S8.**
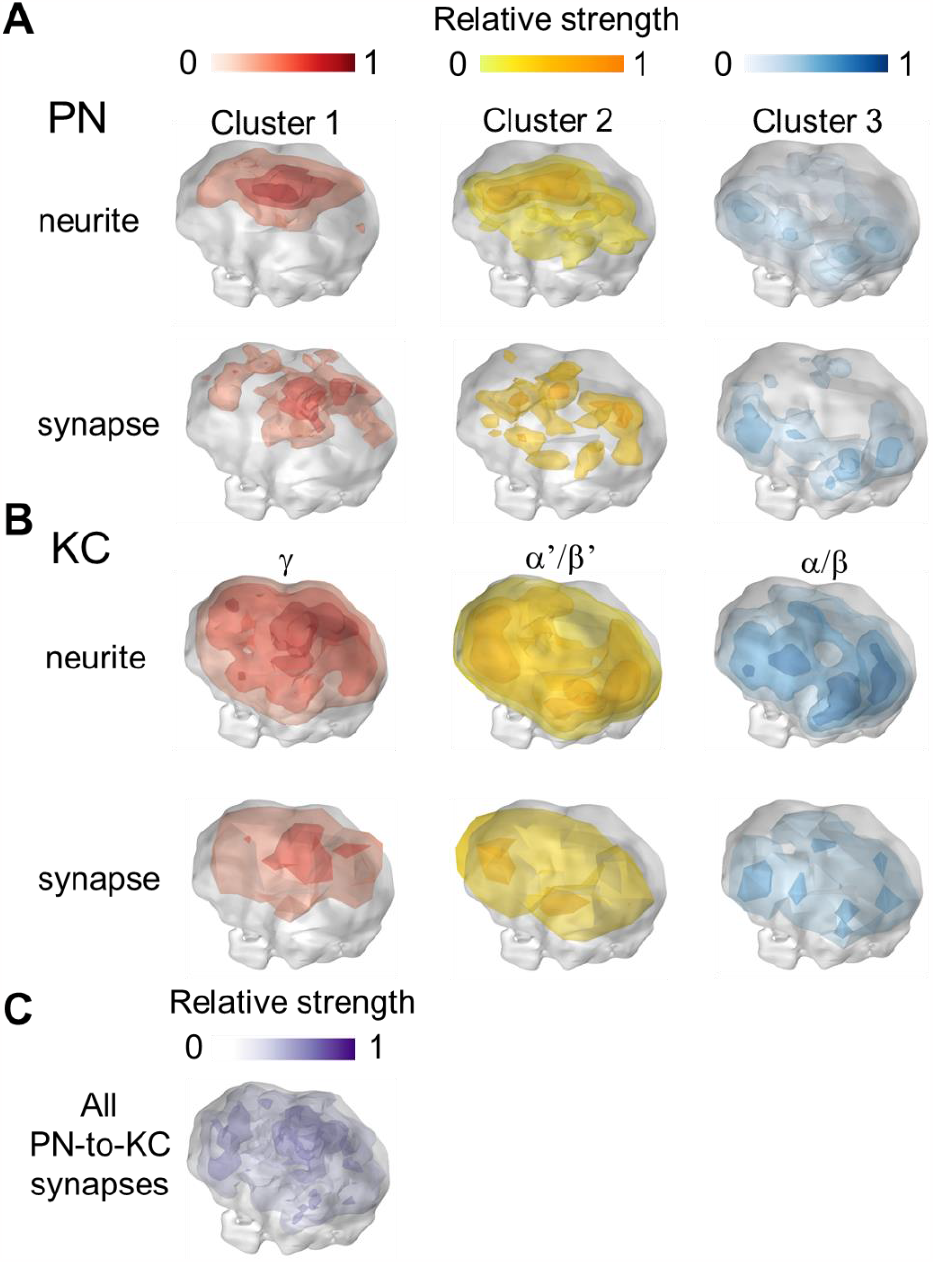
Spatial innervation density of PN clusters and KC classes in CA. (**A**) The density map of PN neurites (top) and PN-to-KC synapses (bottom) for three PN clusters (from left to right). (**B**) same as in (**A**) but for three KC classes. (**C**) The density map of all PN-to-KC synapses.

**Fig. S9.**
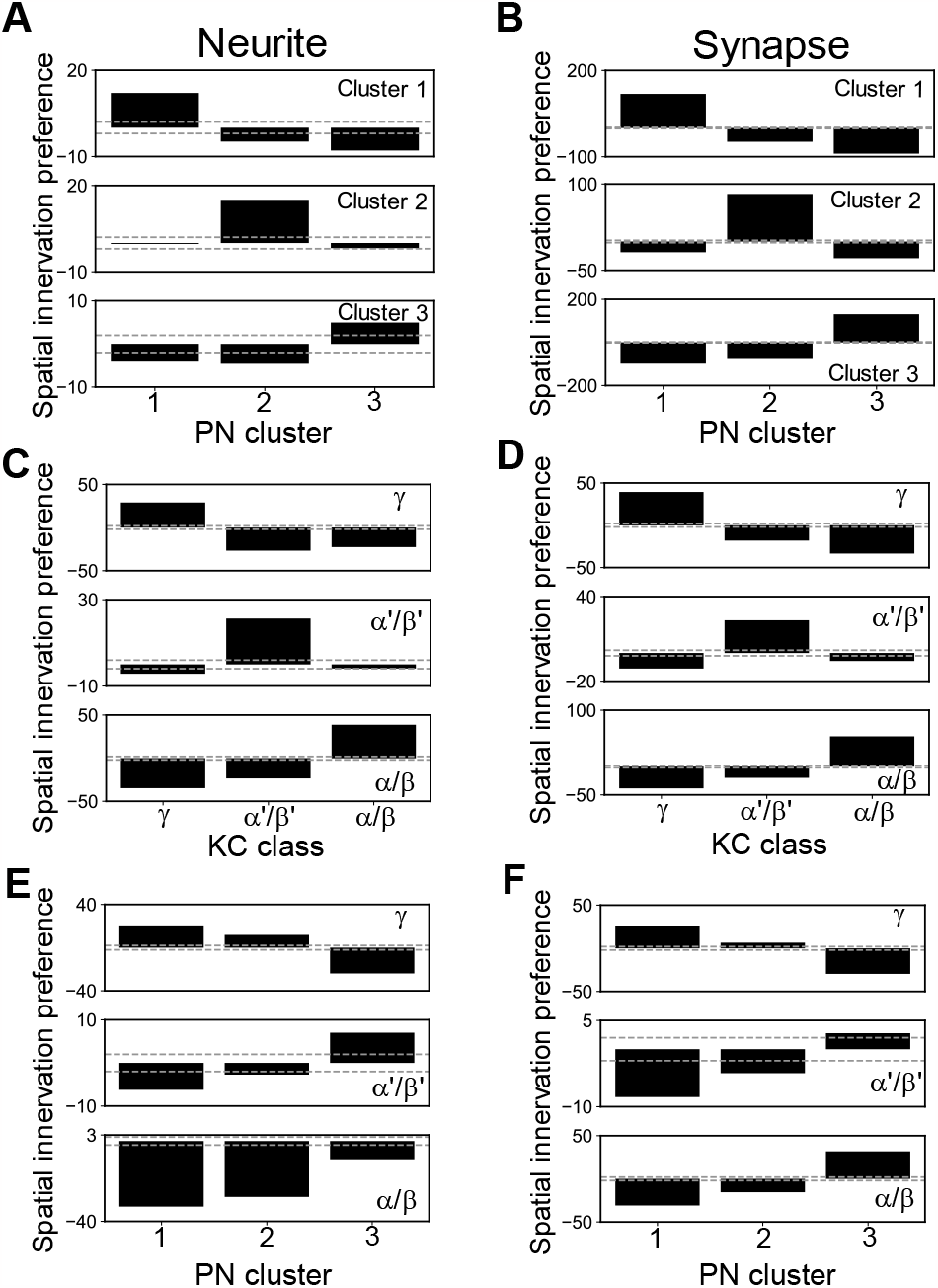
Spatial innervation preference comparison among PN clusters and KC classes. The spatial innervation preference (S) measures the normalized overlap between two sets of structures (neurites of PN clusters 1 and 2 for example) with respect to the shuffled data (see Supplementary Methods). (**A**) Spatial innervation preferences computed based on neurites from the three PN clusters. (**B**) Same as (**A**) but for synapses. (**C**) and (**D**) Same as (**A**) and (**B**) but for three KC classes. (**E**) and (**F**), similar to (**A**) and (**B**) but for the spatial innervation preferences between PN clusters and KC classes. For each plot, the upper and lower dashed lines indicate the preference scores of 2 and -2, respectively.

**Fig. S10.**
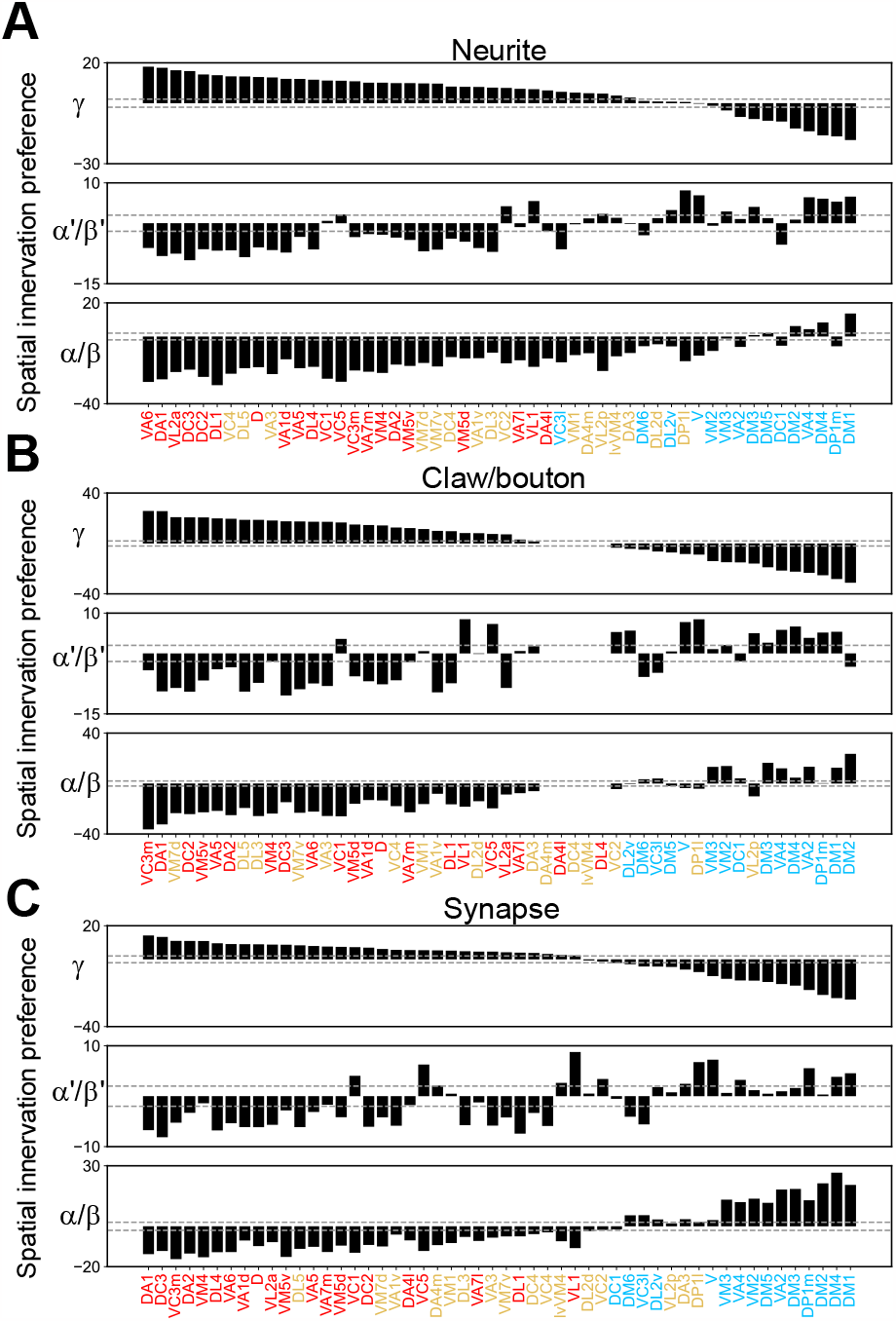
Spatial innervation preference between glomeruli and KC classes. (**A**) Spatial innervation preferences (S, see Supplementary Methods) computed based on neurites. (**B**) same as (**A**) but for claws/boutons. (**C**) Same as (**B**) but for PN-to-KC synapses. The color assignment of each glomerulus is the same as in Fig. 1B. The glomeruli are ordered based on the descending preference score of γ. For each plot, the upper and lower dashed lines indicate the preference scores of 2 and -2, respectively.

**Fig. S11.**
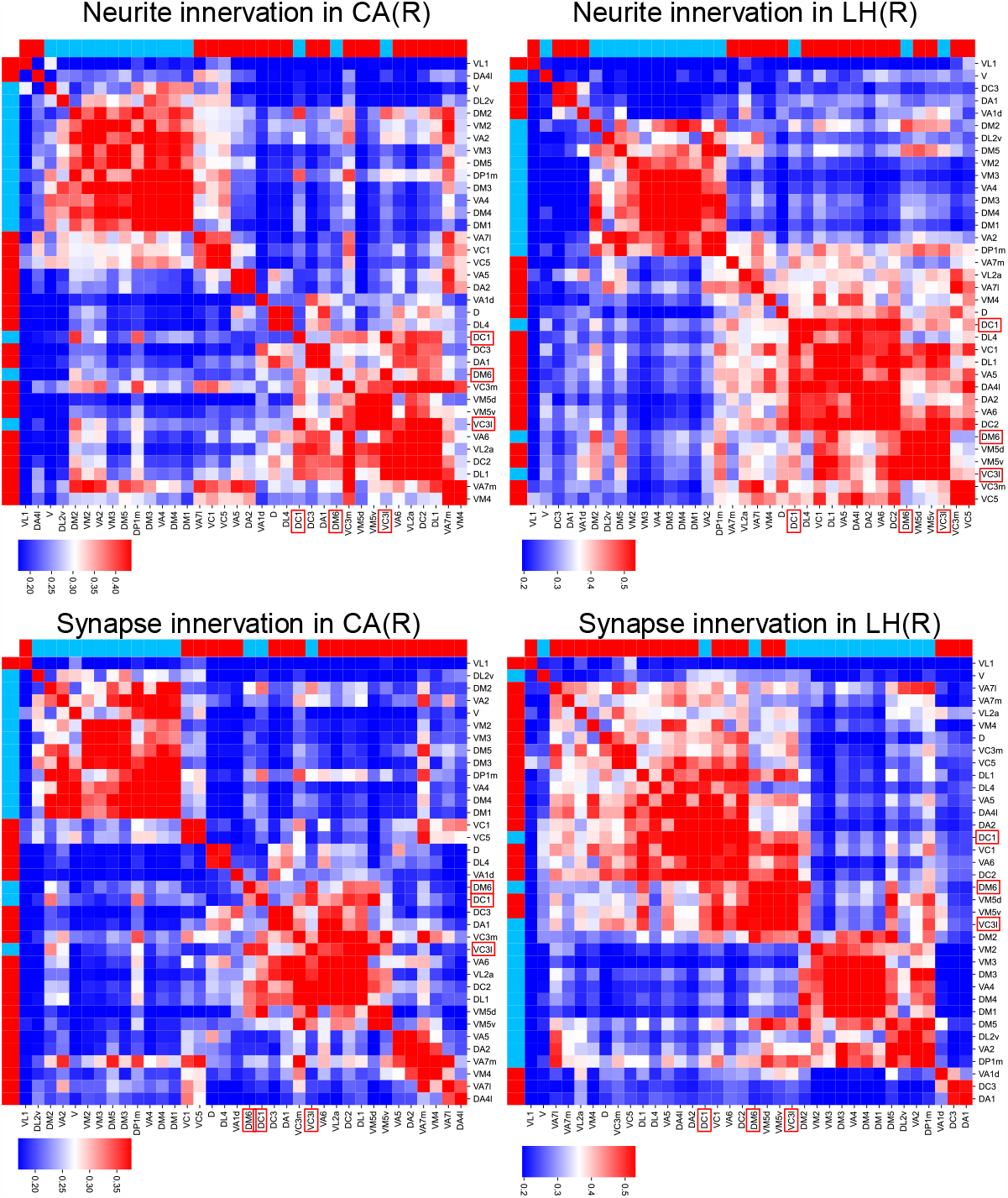
Spatial segregation of PNs in CA and LH. We compute the spatial innervation similarity (see Supplementary Methods) between glomeruli in the right CA (left column) and right LH (left column) for PN clusters 1 and 3. The spatial innervation similarity is computed based on neurites (top row) and synapses (bottom row). The top row and leftmost column in each matrix indicate PN clusters (red for cluster 1 and blue for 3). PN cluster 1 and PN cluster 3 are segregated for most glomeruli in both CA and LH. The spatial innervation pattern of glomeruli, DC1, VC3l, and DM6 (indicated by red frames) in cluster 3, is different from other cluster 3 glomeruli in both CA and LH. The color gradient bar represents the range of spatial innervation similarity values, spanning from the 10^th^ percentile (minimum) to the 90^th^ percentile (maximum) of the similarity distribution among all pairs.

**Fig. S12.**
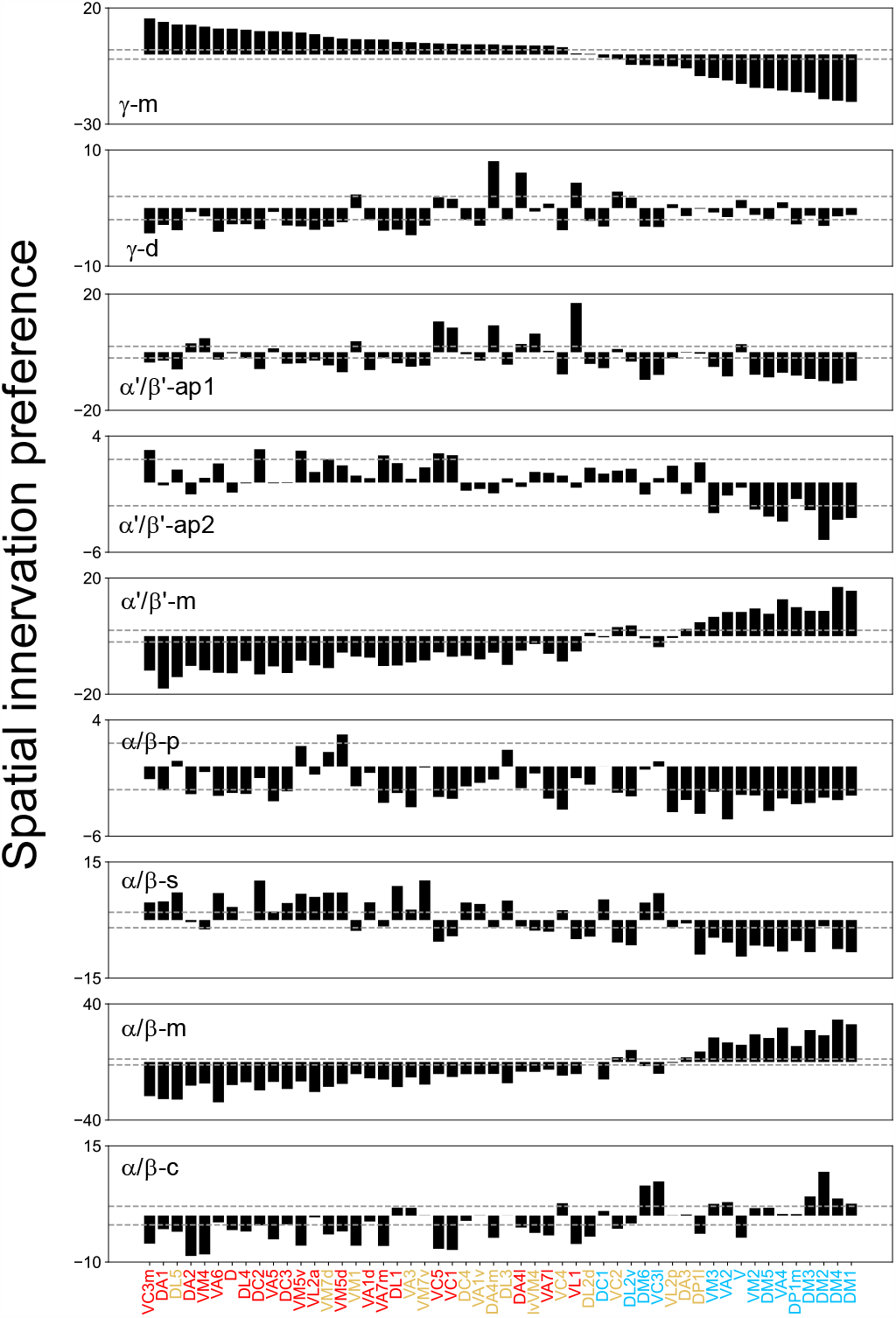
Synaptic spatial innervation preference between glomeruli and KC subclasses. The glomeruli are ordered based on the descending preference score of γ. The glomeruli are colored based on the PN clusters as in Fig. 1B.

**Fig. S13.**
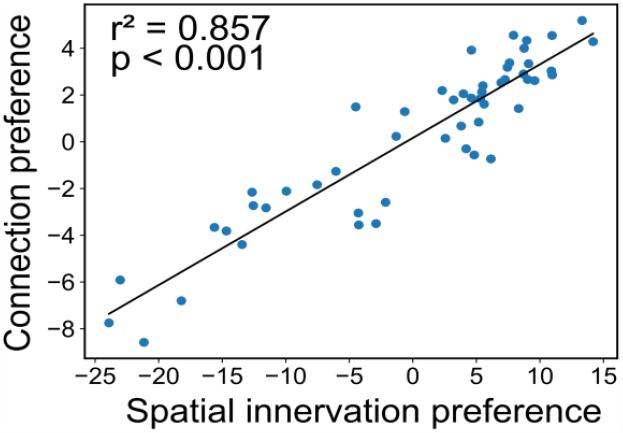
Spatial innervation preference of glomeruli and KC classes is highly consistent with PN-to-KC connection preferences. The connection preference (G_γ_, y-axis in the top panel in Fig. 1B) is strongly correlated with the synaptic spatial innervation preference compared with γ (S_γ_, y-axis in the top panel in fig. S10C).

**Fig. S14.**
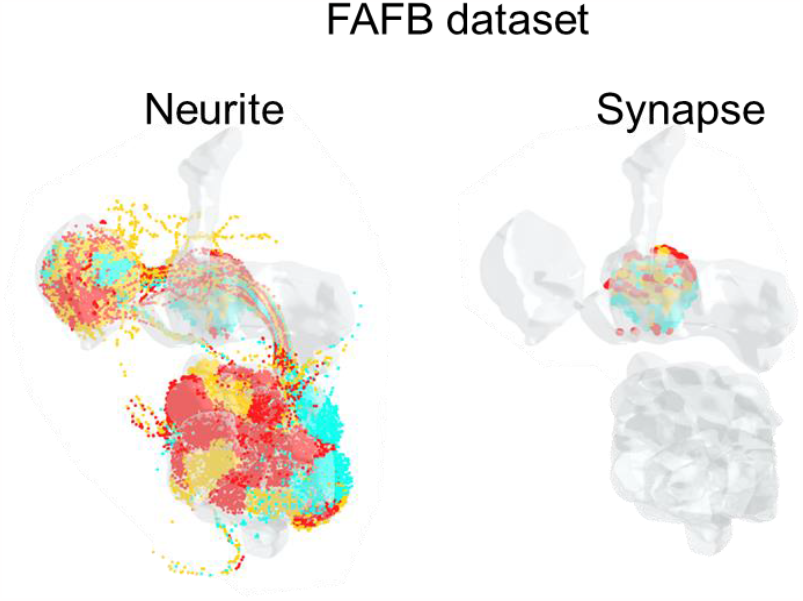
Spatial innervation contour of PN clusters in FAFB dataset. The three PN clusters are represented by three different colors, as in Fig. 1B. The PN cluster 3 projects onto the ventral region of the calyx. Conversely, the PN cluster 1 projects to the dorsal regions of the calyx.

**Fig. S15.**
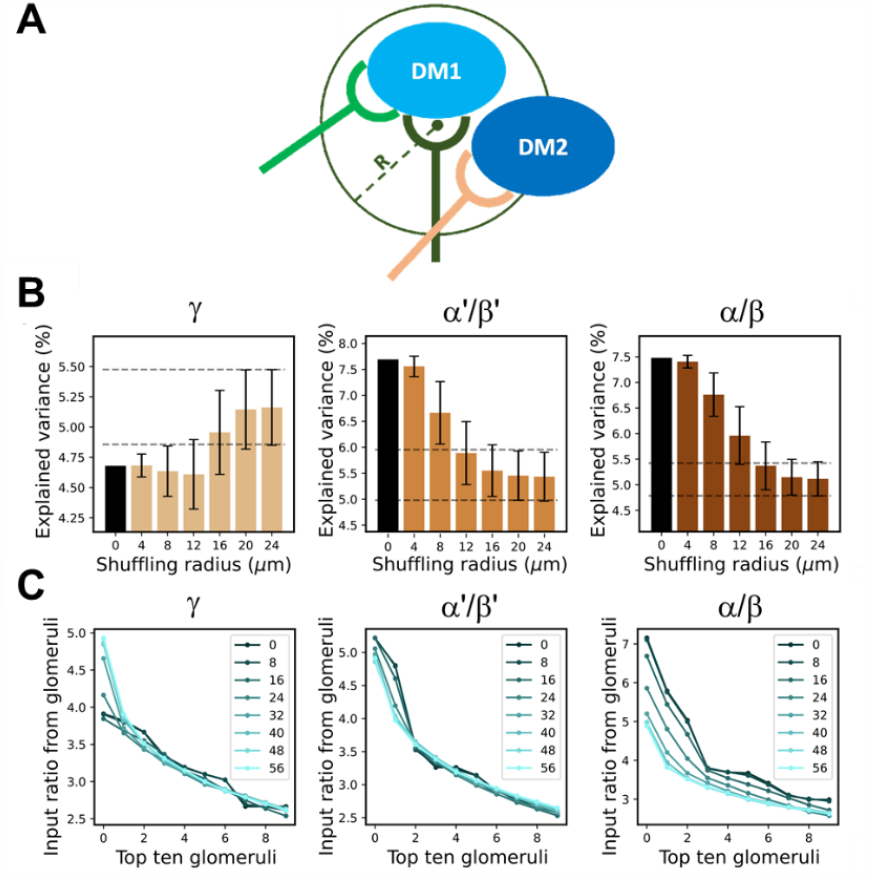
The spatial stereotype of calyx confines the glomerular input diversity of KC classes. (**A**) The schematic plot of the local shuffling algorithm. Two blue ovals illustrate the boutons from different PNs, and the forceps represent the KC claws. The claws with a distance shorter than *R* randomly exchange upstream PNs. The total number of claws and PN connections are maintained during the shuffling. (**B**) The percent variance distribution of each KC class connectivity with local shuffling. (**C**) Input ratio from top ten glomeruli under different local shuffling radii for each KC class.

**Fig. S16.**
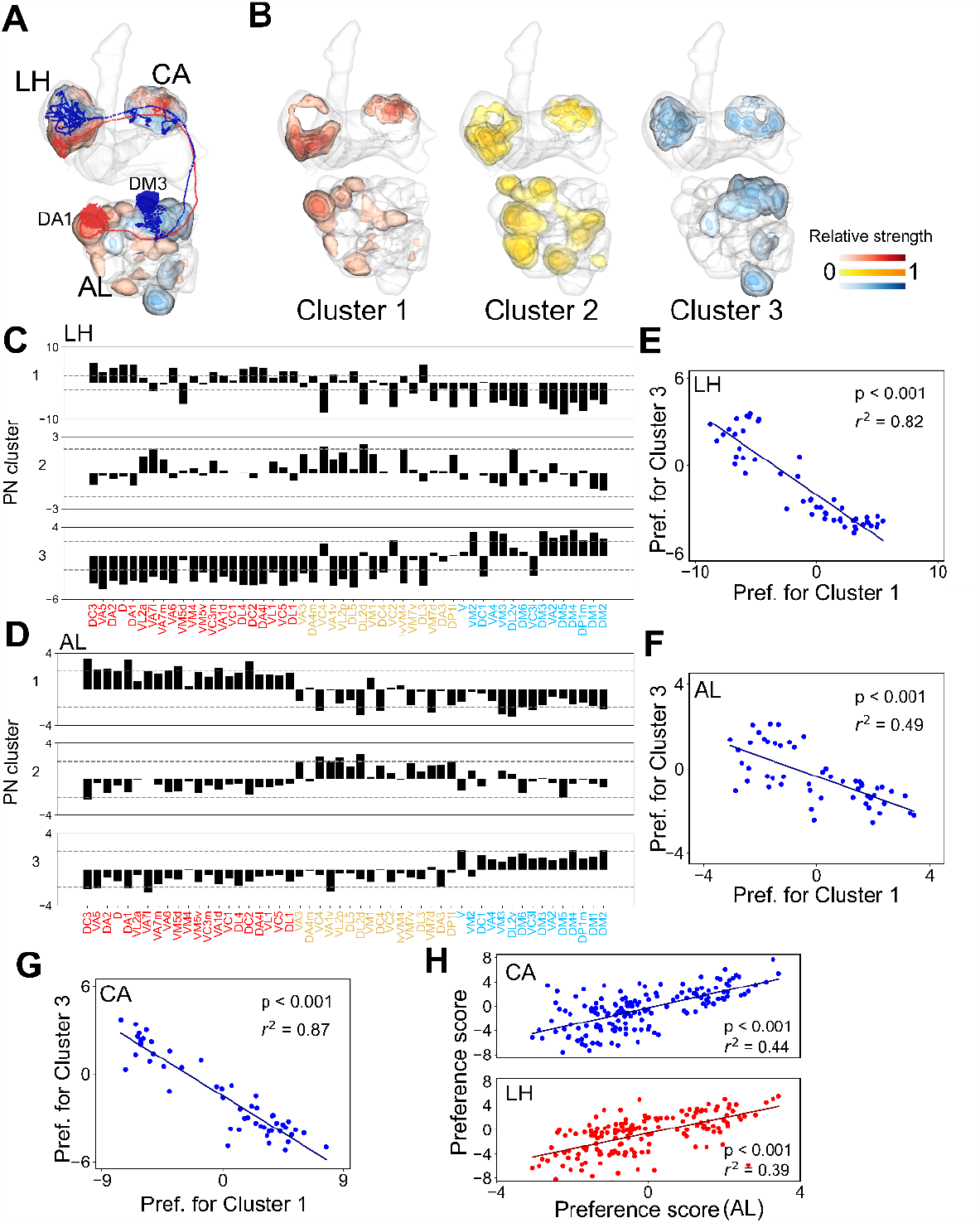
Synaptic spatial segregation of PN clusters across olfactory network architecture. (**A**) The synaptic density maps of Cluster 1 and Cluster 3 PNs with example neurons. (**B**) The synaptic density maps of PN clusters. (**C**) and (**D**) The spatial innervation preference of each glomerulus for three PN clusters. Ordinates indicate the differences in preference between the real and shuffled networks in LH (**C**) and AL (**D**). (**E**-**G**) The regression plot of the preference score of each glomerulus for PN cluster 1 (x-axis) and PN cluster 3 (y-axis) in LH (**E**), AL (**F**), and CA (**G**). (**H**) Correlation of synaptic spatial innervation preference between each glomerulus and each PN cluster (for the comparison between AL and CA, r^2^ = 0.44, p < 0.001; for the comparison between AL and LH, r^2^ = 0.39, p < 0.001).

**Fig. S17.**
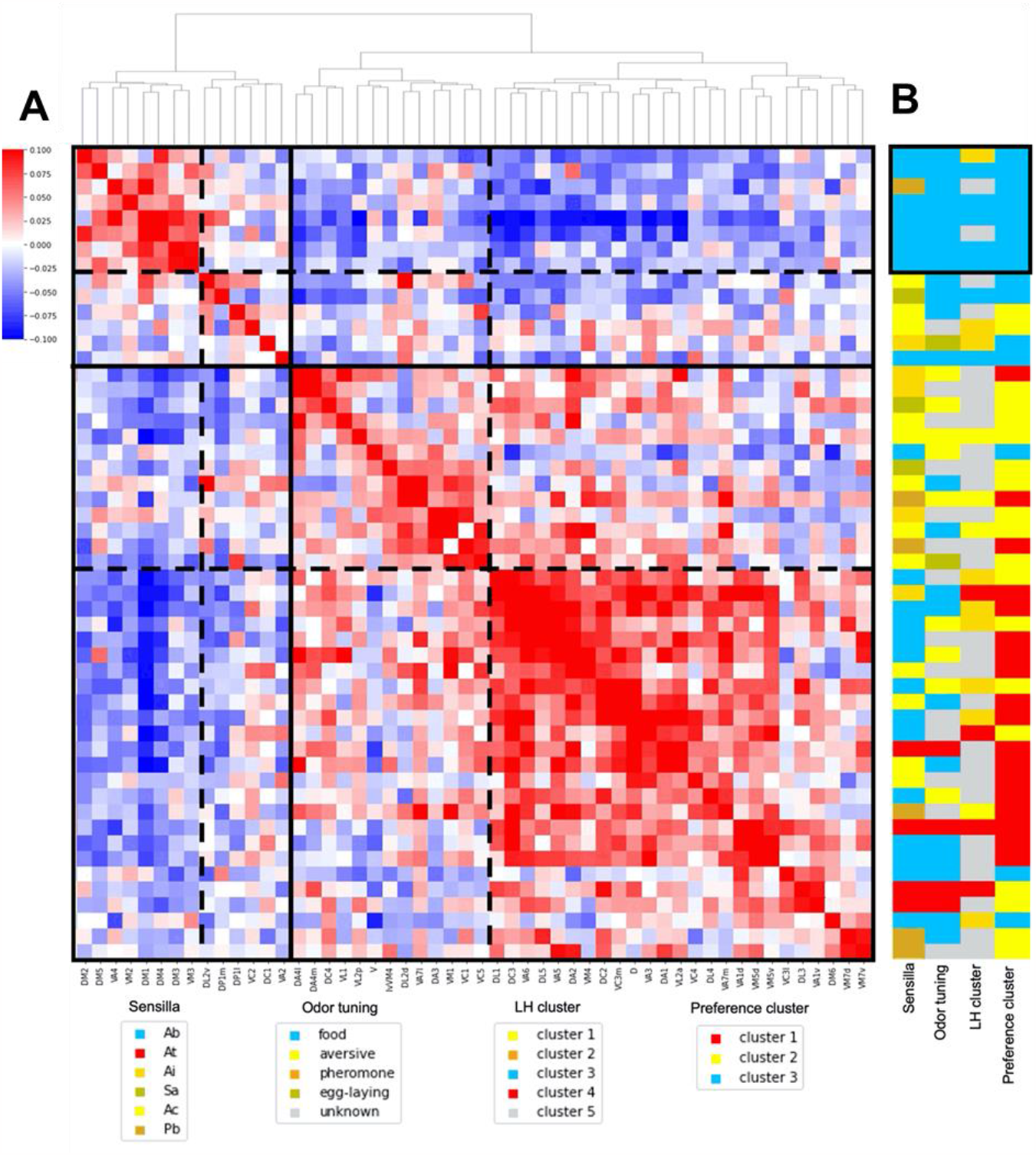
Correlation between projection and physiology for AL glomeruli. (**A**) The correlation matrix of glomerular projection. Color of each element in the matrix represents the similarity (Pearson correlation) of the targeting KCs between glomeruli. Hierarchical clustering analysis is performed to visualize the grouping of glomerular subsets. (**B**) The morphological and functional features of glomeruli are color labeled on the right-hand side of the matrix, including the preference cluster (Fig. 1B), the sensilla type of glomerular upstream ORNs (*38*), and the glomerular axonal cluster in LH (*18*). A group of glomeruli (black rectangle) with similar projection targets shares similar physiological characteristics.

**Fig. S18.**
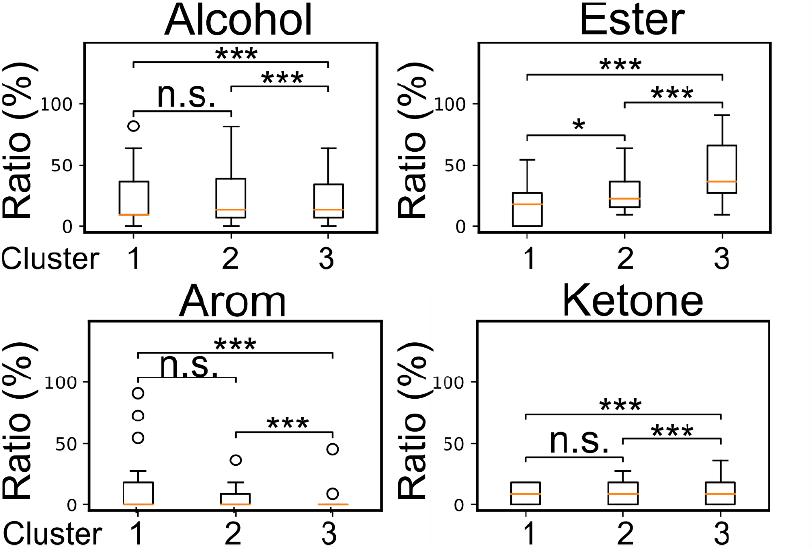
The tuning profile of the glomerular cluster. We counted the ratio of glomeruli that are sensitive to an odor (alcohol, ester, arom, or ketone) in each PN cluster. A glomerulus is defined as sensitive to an odor if it is among the top ten most responsive odors of that glomerulus (data from the DoOR database (*42*)). (The Mann-Whitney U test is used to examine the statistical significance among three glomerular clusters. n.s. for not significant, * for p < 0.05, ** for p < 0.01, and *** for p < 0.001.)

**Fig. S19.**
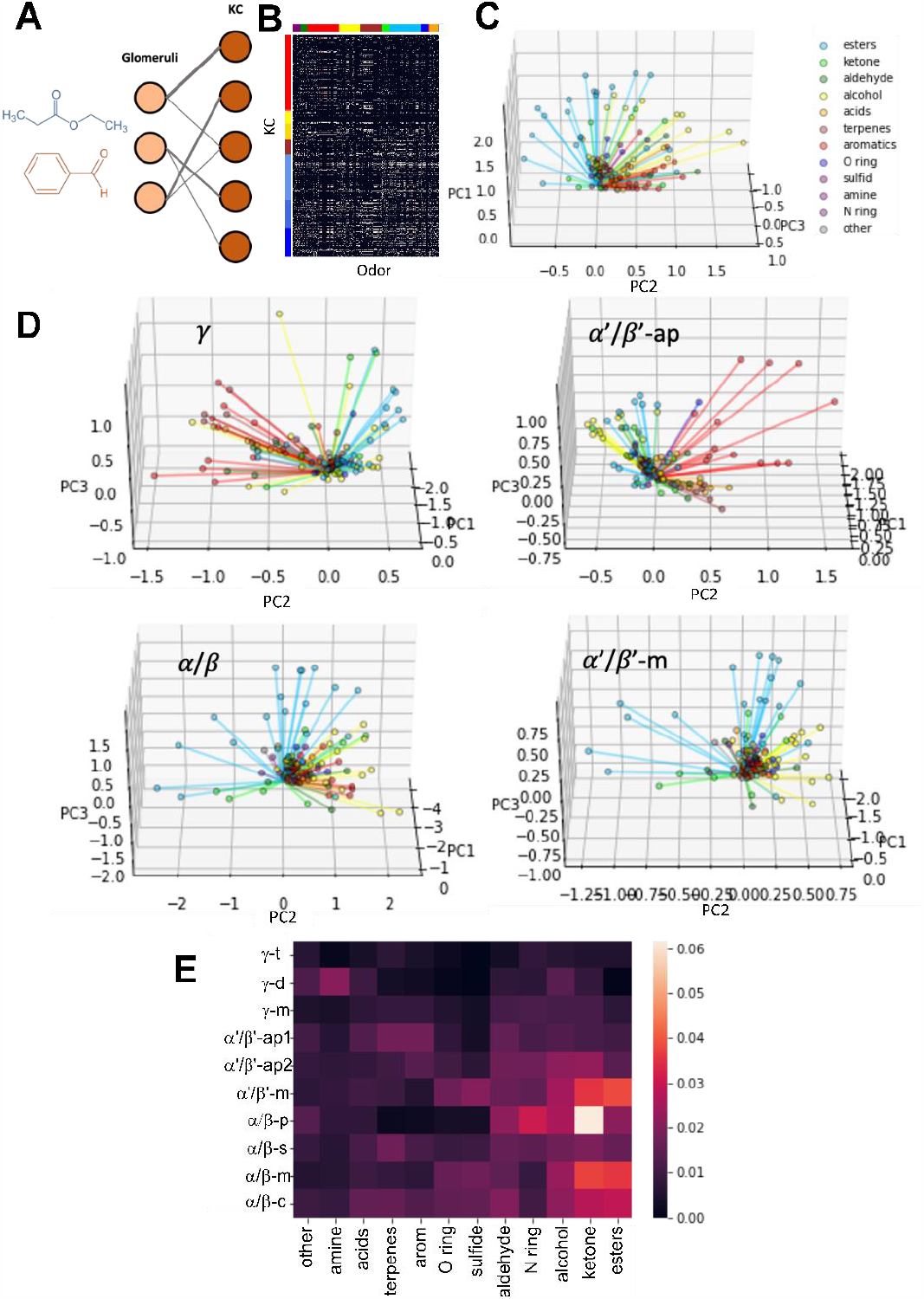
Olfactory response simulation of KCs and MB lobes. (**A**) The schematic illustration of odor representation from AL glomeruli to KCs. The KC response to an odor is simulated by multiplying the glomeruli-to-KC connection number by the glomeruli response level induced by a given odor. The KC response level is normalized by the total claw number of each KC. (**B**) The table of simulated KC response to ∼700 odors. (**C**) The first three principal components in PCA of the simulated odor representation of KCs. Each dot in the plot represents an odor and is color-labeled according to the functional group the odor possesses. (**D**) Similar to (**B**) but for KCs in γ, α’/β’-ap, α’/β’-m, or α/β lobe, respectively. The angular difference between each two molecules represents the KC responsive pattern dissimilarity. (**E**) The chemical sensitivity of KC subclasses is computed by summing the odor responses of KCs in each subclass.

**Fig. S20.**
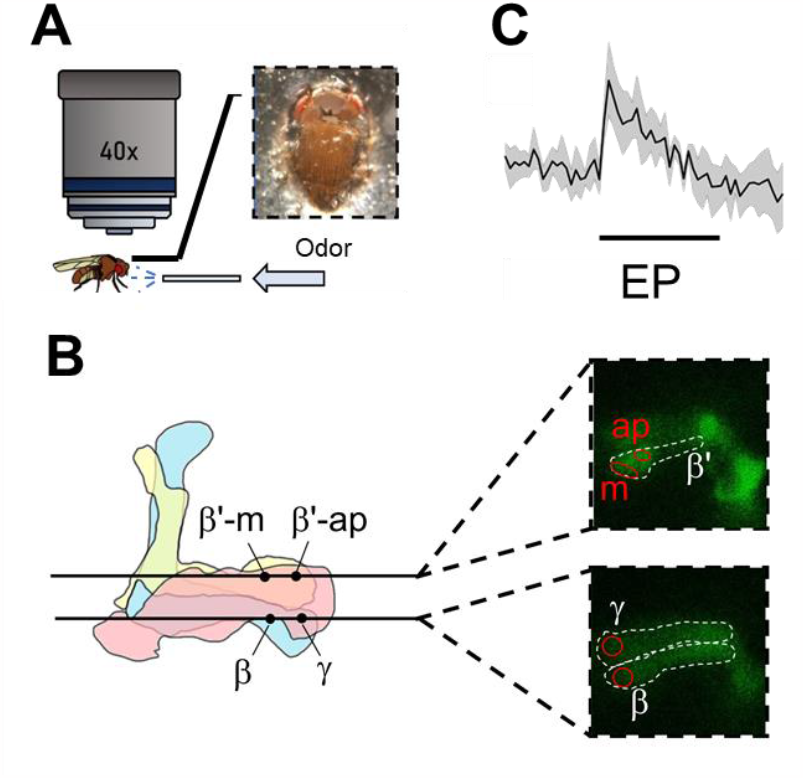
Functional imaging for odor-evoked response in MB lobe. (**A**)The schematics of our functional imaging experiments. (**B**) The recording ROIs in each lobe (β’-ap, β’-m, β, and γ). (**C**) An example calcium response in the β lobe. The bar indicates the period when the odor is delivered.

**Fig. S21.**
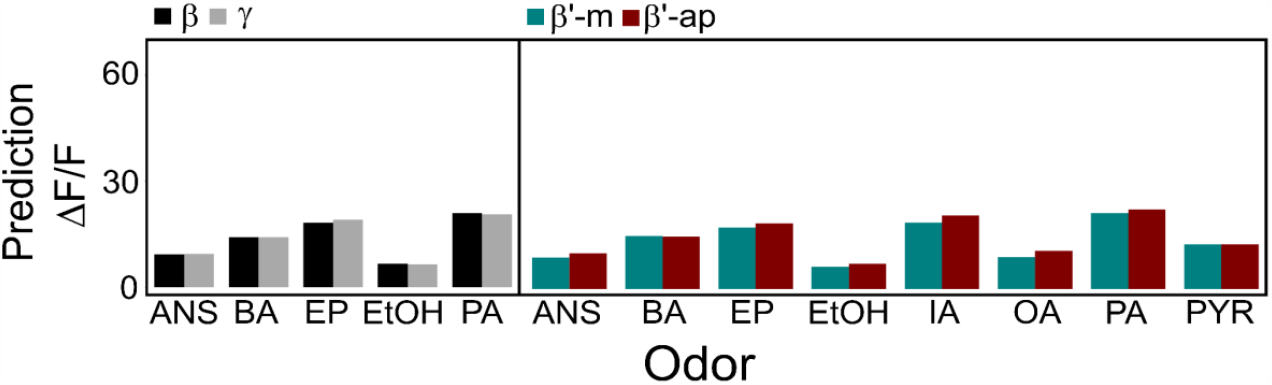
The random networks appease differential odor responses by prediction. The same simulation approach as in Fig. 3C but using shuffled random networks.

**Fig. S22.**
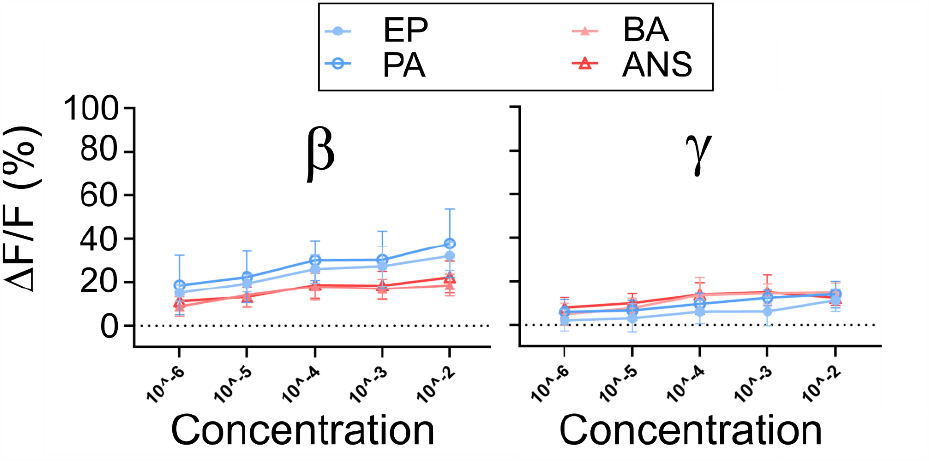
Distinct response levels in different MB lobes induced by esters and aroms. The odor-evoked response levels of MB lobes across six orders of magnitude of odor concentration. Ethyl propionate, EP, and Pentyl acetate, PA, evoke significantly higher responses than BA and ANS in the β lobe but not in the γ lobe.

## Supplementary Methods

### Spatial clustering for glomeruli

We selected glomeruli of PN cluster 1 and PN cluster 3 to perform the clustering analysis using the Python package Seaborn. The color bar indicates the spatial innervation similarity. The color gradient ranges from 10% (minimum) to 90% (maximum) of the similarity values in the matrix. The pairwise distance is defined by Pearson correlation, and the clustering method is ‘*single*’ for better capture similar glomeruli.

### Spatial innervation preference

All neural structures, including the neurites, claws, boutons, and synapses, are represented by spatially distributed points in the data. To estimate the spatial innervation preferences between the neural structures (point sets) A and B, we first calculated the distribution density of the sets of points by kernel density estimation (KDE). Next, we attained the spatial distribution similarity (S) between A and B by calculating the JS divergence (see Methods). Then, we quantified the preference score by

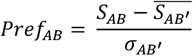

where *S*_*AB*_ is the spatial distribution similarity between A and B. After repeating the categorical shuffling (see Methods) by a number of times, we obtained the mean, 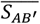, and the standard deviation, *σ*_*AB*′_, of the spatial distribution similarity between A and shuffled B’s. For example, *Pref*_*11*_ represents the spatial innervation preference between the original PN cluster 1 and the shuffled PN cluster 1.

### Global shuffling with synaptic weighting

We aim to incorporate the synapse weighting between PNs and KCs during the shuffling process. The unit of shuffling is a KC claw, and as part of this process, the glomerular PNs are reassigned with new downstream KCs claws. Throughout the shuffling, we ensure the maintenance of the total synapse number for each claw, as well as the total KC claw number connecting with each PN. The synaptic modified G-KC connection table is defined as the sum of the total synapse number between glomerular PNs and their downstream KCs.

**Table S1.** The classification of glomeruli based on the PN projection anatomy in LH (*18, 20*).

|  |  |
| --- | --- |
| **Group 1** | D, DC1, DC2, DL1, DL5, DM2, DM6, VA6, VC2, VL2p, VM7<br>(1 was not collected in the FlyEM dataset.) |
| **Group 2** | DA1, DC3, DL3, VA1d, VA1v, VA3 |
| **Group 3** | DM1, DM3, DM5, DP1m, VA2, VM2, VM3<br>(DL6 was not collected in the FlyEM dataset.) |
| **Group 4** | VA7l, VA7m, VL2p, VM1 |
| **Group 5** | DA1, DA3, VA1v, VL1 |

**Table S2.** The classification of glomeruli based on the PN projection anatomy in MB (*18, 20*).

|  |  |
| --- | --- |
| **Group 1** | DM2, DM5, DP1m, VA7m, VL2p, VM2, VM3 |
| **Group 2** | VA1d, VA7l, VC2, VL2p |
| **Group 3** | DA1, DL3, DL6, VM7 (DL6 was not collected in the FlyEM dataset.) |
| **Group 4** | 1, D, DC2, DC3, DL1, DM6, VA1v (1 was not collected in the FlyEM dataset.) |

**Table S3.** The classification of glomeruli based on their sensilla (*38*).

|  |  |
| --- | --- |
| **LB (ab)** | DM1, DM2, DM4, DM5, V, VA2 |
| **TB (ab)** | D, DL4, DM6, VA3, VC4, VM2, VM3, VM5<sub>v</sub> |
| **SB (ab)** | DA2, VA6, VA5, DM3, DL5, DL1 |
| **pb** | 1, VA4, VA7<sub>l</sub>, VC1, VC2, VM7 (1 was not collected in the FlyEM dataset.) |
| **ac** | DC4, DL2, VC3, VL1, VL2<sub>a</sub>, VM1, VM4, VM6 (DL2 and VC3 have two subgroups in FlyEM. VM6 was not collected in the FlyEM dataset.) |
| **T3** | DA4<sub>l</sub>, DA4<sub>m</sub>, DC1, DL3, VA1<sub>d</sub>, VA1<sub>v</sub> |
| **T2** | DA3, DC3 |
| **T1** | DA1 |

**Table S4.**
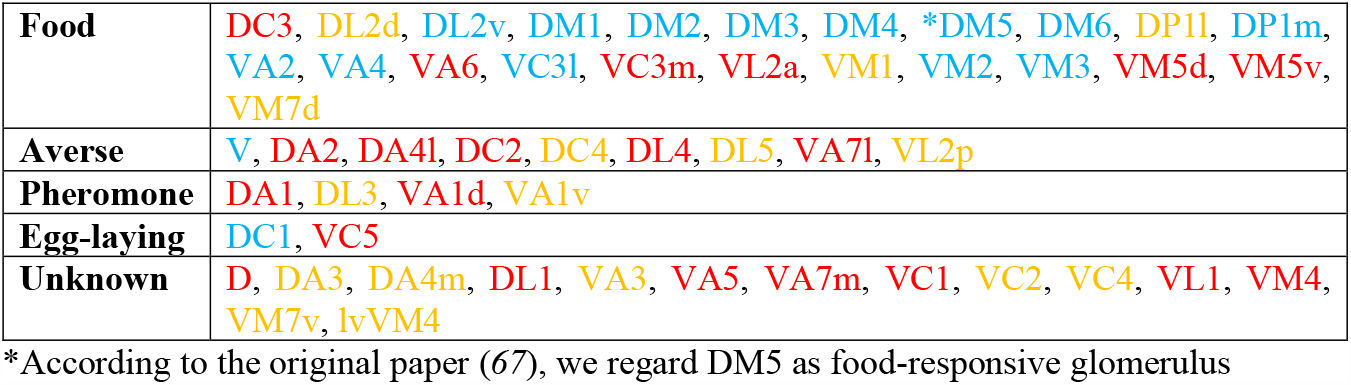
The classification of glomeruli based on their functionality (*22*).

**Table S5.** The community structure derived from the FAFB dataset (*22*).

|  |  |
| --- | --- |
| **Core** | DM2, DP1m, VM2, DL2v, DM3, DM4, DM1, VM3, VA2, VA4 |
| **2<sup>nd</sup>** | DM6, DM5, DC1, DL2d, VC4, VC3l, DP1l, VA6, VM5d |
| **Under-convergent** | VA7m, VP1, VP2, V, VL1, DA4m, DA2, DA3, DA1<br>(VP1 and VP2 were not considered in our analysis) |
| **Others** | DL1, DC3, VA7l, VL2a, DC2, VM1, D, VM4, VA1d, VM7v, VA5, VC3m, VC1, DL3, DL5, VA3, DC4, VM7d, VC5, VA1v, DA4l, VL2p, DL4, VM5v, VC2, VP3 (VP3 was not considered in our analysis) |

**Table S6.**
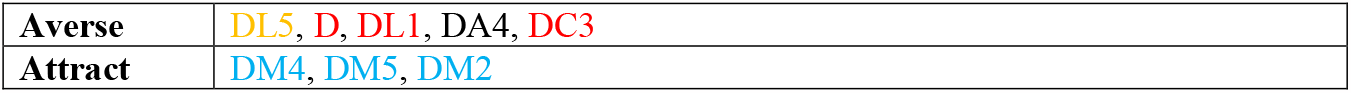
The glomeruli associated with aversive or attractive responses based on 12 aversive and attractive odors (6*8*).

**Table S7.** The classification of glomeruli based on the innervation overlap between PN and KC (*17*).

|  |  |
| --- | --- |
| $\gamma$ | DL1, VL2a, VA5, VA7l, DC2 |
| $\alpha'/\beta'$ | DM1, VA4 |
| $\alpha/\beta$-p | DL4 |
| $\alpha/\beta$-s | DL3, DA1, VA1d, D |
| $\alpha/\beta$-c | DM2 |

**Table S8.** The classification of glomeruli based on the neuronal distance (*30*).

|  |  |
| --- | --- |
| **1** | DL5, VP4, VP1l |
| **2** | VM4, VC5, VM5d, VM6, DL5, VC3, DC4, VP5 |
| **3** | DL2v, DL2d |
| **4** | VM4, VL1, VP2 |
| **5** | VC5, DA4m, V, VP4, VP2, VP1d, VP1m |
| **6** | VM1, VL1, DM5, VA4, VC2, DA3, VA7m, DP1l, DP1m, VL2p, DA2, DA4l, VC1, VA5, VA7l, DA1, VP2 |
| **7** | DL2v, DM5, DM1, DM4, VM2, DM2, DA2, DM3 |
| **8** | VL2a, DC1, D, DL4, VA1v, DA1, DC3, VA1d |
| **9** | VM1, VM5d, VA3, DA3, DL1, VM7v, VC4, VA2, DA2, D, DC2, VA6, DL3 |
| **10** | DM6, VM5d, VM5v, VM7v, VC3, DL3 |

**Table S9.** Glomeruli and associated receptors with the most responded odor (from the DoOR database (*42*)). The glomeruli are arranged in the hierarchical order, shown in Fig. 3.

|  |  |  |  |  |  |
| --- | --- | --- | --- | --- | --- |
| DM2 | Or22a | ethyl hexanoate | VC5 | lr75d | pyrrolidine |
| DM5 | Or85a | ethyl 3-hydroxybutyrate | DL1 | Or10a | methyl salicylate |
| VA4 | Or85d | ethyl pentanoate | DC3 | Or83c | farnesol |
| VM2 | Or43b | ethyl trans-2-butanoate | VA6 | Or82a | geranyl acetate |
| DM1 | Or42b | ethyl acetate | DL5 | Or7a | E2-hexenal |
| DM4 | Or59b | methyl acetate | VA5 | Or49b | 2-methylphenol |
| DM3 | Or47a | pentyl acetate | DA2 | Or56a | nan |
| VM3 | Or9a | 3-hydroxy-2-butanone | VM4 | lr76a | nan |
| DL2v | lr75abc | acetic acid | DC2 | Or13a | 1-octen-3-ol |
| DP1m | lr64a | 2,3-butanediol | VC3m | lr41a | putrescine |
| DP1l | lr75abc | acetic acid | D | Or69a | ethyl 3-hydroxy hexanoate |
| VC2 | Or35a | 1-heptanol | VA3 | Or67b | acetophenone |
| DC1 | Or19a | valencene | DA1 | Or67d | nan |
| VA2 | Or92a | 2,3-butanedione | VL2a | lr84a | phenylacetaldehyde |
| DA4l | Or43a | 1-hexanol | VC4 | lr75d | pyrrolidine |
| DA4m | Or2a | 3-hydroxy-2-butanone | DL4 | Or49a | 2-heptanone |
| DC4 | lr64a | acetic acid | VA7m | Or33c | cyclohexanone |
| VL1 | lr75d | pyrrolidine | VA1d | Or88a | pyrrolidine |
| VL2p | lr31a | 2-oxovaleric acid | VM5d | Or85b | butyl acetate |
| V | Or21a | carbon dioxide | VM5v | Or98a | ethyl benzoate |
| lvVM4 | lr40a | nan | VC3l | Or67c | ethyl lactate |
| DL2d | lr75abc | acetic acid | DL3 | Or65a | pyrrolidine |
| VA7l | Or46aA | 4-methylphenol | VA1v | Or47b | (S)-(+)-carvone |
| DA3 | Or23a | 1-pentanol | DM6 | Or67a | butyl propanoate |
| VM1 | lr92a | dimethylamine | VM7d | Or42a | propyl acetate |
| VC1 | Or71a | 4-ethyl guaiacol | VM7v | Or59c | ethyl butyrate |

